# Long-term unsupervised recalibration of cursor BCIs

**DOI:** 10.1101/2023.02.03.527022

**Authors:** Guy H. Wilson, Francis R. Willett, Elias A. Stein, Foram Kamdar, Donald T. Avansino, Leigh R. Hochberg, Krishna V. Shenoy, Shaul Druckmann, Jaimie M. Henderson

## Abstract

Intracortical brain-computer interfaces (iBCIs) require frequent recalibration to maintain robust performance due to changes in neural activity that accumulate over time. Compensating for this nonstationarity would enable consistently high performance without the need for supervised recalibration periods, where users cannot engage in free use of their device. Here we introduce a hidden Markov model (HMM) to infer what targets users are moving toward during iBCI use. We then retrain the system using these inferred targets, enabling unsupervised adaptation to changing neural activity. Our approach outperforms the state of the art in large-scale, closed-loop simulations over two months and in closed-loop with a human iBCI user over one month. Leveraging an offline dataset spanning five years of iBCI recordings, we further show how recently proposed data distribution-matching approaches to recalibration fail over long time scales; only target-inference methods appear capable of enabling long-term unsupervised recalibration. Our results demonstrate how task structure can be used to bootstrap a noisy decoder into a highly-performant one, thereby overcoming one of the major barriers to clinically translating BCIs.

## Background

Brain-computer interfaces (BCIs) can enable people with paralysis to interact with computers by directly translating neural activity into command signals. Noninvasive methods such as MEG/EEG and fMRI allow minimal-risk access to brain signals but are limited by low spatial and temporal resolution, respectively ^1,2^ By contrast, intracortical BCIs (iBCIs) benefit from unparalleled spatial and temporal resolution recordings of local neuronal ensembles, resulting in some of the highest-performance communication systems to date ^3,4^.

Despite these advances, intracortical BCIs often require daily recalibration to maintain high performance in the face of signal nonstationarities ^3,5,6^. Neural features exhibit drift over time, reflecting (among other causes) array movements ^7^, device degradation ^8^, physiological changes in single neurons ^9–11^, and the influence of varying behavior ^12,13^. This drift operates on multiple timescales. Single and multi-unit baseline firing rates can change even after a few minutes ^5,9^ while the selectivity of their responses often vary on the order of days (^12^, but see ^14^). As changes compound, decoders fit to a particular time period progressively worsen, resulting in a need for repeated recalibration. Recent efforts at building intrinsically robust decoders ^15,16^ have shown promise counteracting these changes, but nevertheless only delay the time needed until recalibration is needed. Standard recalibration procedures require ground-truth target labels for supervised training; consequently, an iBCI user has to carry out a predefined sequence of training examples and cannot engage in free use of their device during these times.

An alternative approach that has been explored is the use of electrocorticography (ECoG) signals, which involves recording electrical activity from the surface of the brain with coarser electrode grids^2^. Owing to the larger electrode sizing, ECoG signals tend to have a robustness advantage over intracortical recordings ^17–19^ and several studies have demonstrated the potential of ECoG for longer-term use in BCIs. For example, Silversmith and colleagues ^17^ used ECoG signals to enable a patient with brainstem stroke to control a cursor on a computer screen over roughly one month. However, ECoG signals do not have the same spatial resolution as intracortical recordings, with each electrode recording the summed activity of many thousands of neurons^20^, and therefore result in lower-bandwidth cursor BCI control ^3,21,22^.

To maintain the benefits of high-SNR intracortical recordings while mitigating signal instability, many groups have examined unsupervised recalibration methods for decoders ^5,6,23–28^. These are algorithms that recalibrate using the neural features only and do not require ground-truth knowledge of where the targets are located, which one was cued, or even the user’s intentions. Recent work has focused on domain mapping strategies ^6,25,27–29^, where a function *f* is sought such that the distribution of neural features from an unlabeled test set *P_test_* matches those from a training period *P_train_* when mapped through f, i.e. *f*(*P_test_*) ~ *P_train_*. In this case, the assumption is that with proper choice of *f*, the subsequent mapping from features to targets is preserved. For noisy high-dimensional neural recordings, *f* usually operates in a low-dimensional subspace that encodes a majority of task-related modulation ^6,11,30,31^. In particular, factor analysis (FA) stabilization ^6^ uses a FA model to identify task subspaces on two different days. A Procrustes realignment within these spaces is then used to realign new data so that old decoders, trained on the old subspace, can work. More recently, Farschian and colleagues ^25^ improved upon FA stabilization by leveraging deep learning for nonlinear alignment (“adversarial domain adaptation network”, or ADAN).

In this work, we propose an alternative approach. Instead of stabilizing the feature distribution P(x) through a domain mapping strategy, we make use of P(y), which is the prior knowledge of task structure, by leveraging a probabilistic model to infer user intentions during 2D cursor control. This is analogous to how language models denoise earlier stages of automatic speech recognition systems, making use of linguistic structure to guide inferred speech estimates. Our method provides inferred target labels (“pseudo-labels”) at each timestep, alongside confidence estimates, that are used to recalibrate the cursor decoder using standard optimization approaches. This strategy allows for repeated recalibrations across time, enabling adaptation to accumulating neural changes that might otherwise prevent long-term decoder use. As we model target information using decoder outputs (instead of neural signals or latent representations in a decoding pipeline), our approach is agnostic to the choice of decoder or input signal. This approach is similar to the self-training strategies explored in prior BCI work ^24,26^, where a decoder’s outputs are used to correct itself. Here we extend the framework from discrete classification ^24^ and non-human primates (NHPs) ^24,26^ to a continuous regression setting with a human user. Our model is inspired by retrospective target inference (RTI) ^5,32^, a method that infers targets from user selections to then recalibrate velocity decoders. Here, we build on this intuition by leveraging cursor behavior in a structured, probabilistic framework that enables confidence estimates associated with model predictions across all timesteps using a Hidden Markov Model (HMM). We hence refer to this approach as “PRI-T” (Probabilistic Retrospective Inference of Targets).

Since PRI-T models target information using decoder outputs (instead of leveraging latent representations in neural data), it doesn’t require a dimensionality bottleneck in decoder architectures that risks tossing out task-relevant information (this is likely ameliorated however by more powerful dimensionality reduction techniques e.g. ^27,28,33^) and has the potential for arbitrary retraining of multi-layer models. By contrast, neural latent space methods currently update only early components of a decoder (e.g. a single layer in FA stabilization, an autoencoder network with ADAN, and an alignment network in NoMAD, ^28^). Finally, PRI-T is signal-agnostic: since it uses structure in cursor outputs it can theoretically be deployed with BCIs that leverage other control signals such as local field potentials ^34,35^, ECoG ^17,36^, and EEG ^37^.

To develop the approach, we used 73 recording sessions spanning five years from a participant (referred to as ‘T5’) enrolled in the BrainGate2 pilot clinical trial. In the Results, we show that PRI-T, FA stabilization ^6^, and ADAN ^25^ all perform equivalently when applied offline to pairs of days, but with a crucial caveat: All methods fail when day pairs are separated by large amounts of time. We further study this in large-scale simulations ^38^, and demonstrate that unsupervised recalibration procedures require iterative or “chained” applications across subsequent sessions to maintain robust performance across time. Despite modifying FA stabilization for iterative deployment, we nevertheless find that only target labeling strategies are able to effectively deliver long-term control in simulation; domain mapping with FA stabilization appears to accumulate error until eventually diverging. Hence, PRI-T appears similar to existing approaches in pairwise recalibrations but outperforms in iterative, longer-term settings. Finally, we use PRI-T to obtain one month of closed-loop control in participant T5 without any supervised retraining.

## Results

### Characterization of nonstationarities across 5 years

We first analyzed 73 closed-loop cursor control sessions from T5 to characterize the magnitude and time scale of the neural nonstationarities present in the recordings and their potential impact on decoding performance. These results set the stage for the rest of this work and inform the design of methods that aim to accommodate these nonstationarities.

During these sessions, T5 attempted to move a computer cursor to a series of targets while neural activity was recorded from two 96-channel silicon microelectrodes (Blackrock Neurotech) implanted in “hand knob” area of dorsal motor cortex (Fig. 1A). A linear decoder translates the neural activity into cursor velocity signals, thus enabling control of 2D graphical user interfaces with a computer mouse. Much like a mouse enables able-bodied users to issue discrete clicks on parts of a screen, the BCI system also enables selections through neurally-derived ‘clicks’, which are inferred from a separate decoder, or through a “dwell”-based approach where items are selected if the mouse hovers for sufficient time over them. In these data, both approaches for selecting buttons (‘targets’) are present.

**Figure 1.**
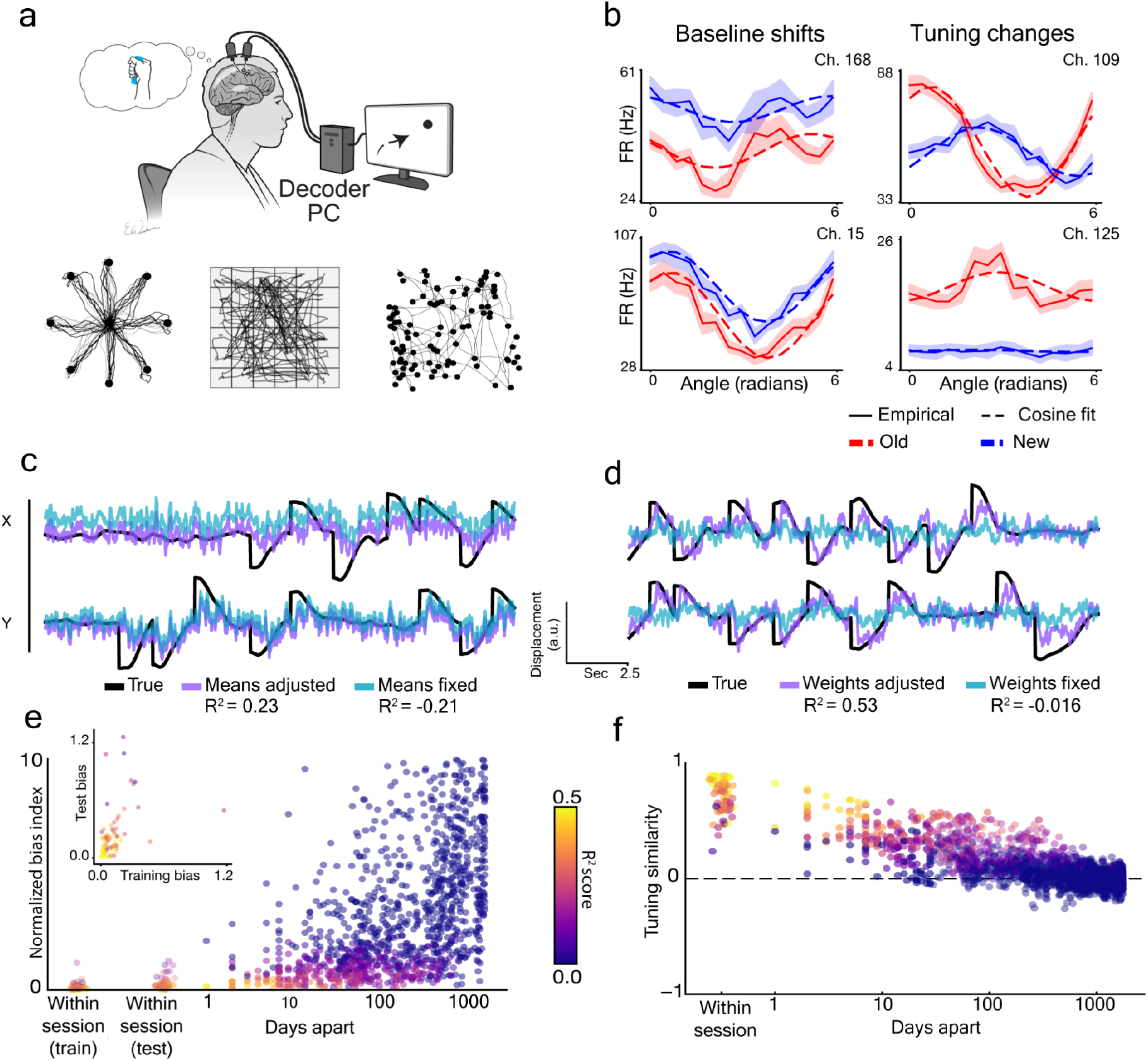
Neural nonstationarities and their impact on cursor control. **(a)** During BCI cursor control, participant T5 attempted to move his hand as if controlling a joystick while recorded neural activity was used to drive a velocity decoder. Cursor control was assessed with different target configurations, such as radial-8 reaches, grids, and random target locations (left to right insets, respectively). Cursor trajectories are plotted for random trials from each target configuration. **(b)** Example neural nonstationarities from two distinct sessions (2 days apart). Dotted lines correspond to cosine tuning models fit within each session solid lines are empirical firing rates for different angles with respect to target position (shading denotes bootstrapped empirical 95% CI). **(c)** Comparison of an offline velocity decoder with and without adaptive mean subtraction. Miscalibrated mean estimates result in low performance on held out data (*R^2^* = −0.21). Predictions are still correlated with ground-truth, implying that weak performance is due to a strong bias rather than tuning mismatch. After adjusting for mean shifts, performance rises to *R^2^* = 0.23. **(d)** Performance when neural tuning has changed greatly. An old decoder fails to work (*R*^2^ = −0.016) despite mean updates, while a new decoder works well (blue, *R*^2^ = 0.53). **(e)** Bias as a function of time between decoder fitting and prediction, plotted on a log-log scale. Each dot corresponds to a pair of sessions, where a decoder is trained on the first and deployed on a second; the new session’s data are centered using the mean estimates from the last block of the old session. Points are colored by corresponding *R*^2^ values. On the left, we also include bias estimates within-session when using adaptive decoders on both training and holdout blocks. Bias estimates tended to rise in holdout blocks despite bias mitigation (inset; p < 2e-8, Wilcoxon signed-rank test). **(f)** Cosine of the angle between decoder tuning weights across pairs of sessions. Values of 1 indicate complete overlap (up to scaling) whereas 0 indicates orthogonal weights. Points are colored by offline *R*^2^ on new sessions after mean recalibration.

We first investigated two classes of neural nonstationarity: baseline shifts and preferred direction (PD) changes under cosine tuning (which captures linear tuning to the user’s intended movement direction, see Methods; Fig. 1B) Baseline shifts maintain a channel’s response specificity to direction but alter its mean activity, whereas PD changes are changes in the preferred direction angle or modulation strength of the tuning curve. Functionally, their impacts on decoder performance also differ. Mean shifts preserve correlations between the predicted and ground-truth velocity with a linear decoder when evaluated offline (see, for example, Perge and colleagues ^9^ for a discussion of closed-loop effects). In an example dataset, we show how a decoder with miscalibrated mean estimates performs poorly on newer data (Fig. 1C light blue traces, “Means fixed”). After adjusting for the bias via updated means, performance rises (“Means adjusted”). By contrast, PD changes cause the decoded predictions to become decorrelated from ground truth (Fig. 1D). Using a ‘stale’ decoder (i.e. a decoder from a previous day, potentially months or even years in the past) with updated means fails to save performance in the face of PD changes (Fig. 1D, “Weights fixed”) whereas performance after supervised recalibration is much higher (“Weights adjusted”).

We examined the timecourses of these changes using two functional metrics that reflect decoder performance: a “normalized bias index” and signal-to-noise ratio (SNR) measure (see Methods). The normalized bias index is a ratio of the bias vector magnitude compared to the decoded control signal’s average magnitude when far from the target (when BCI users engage in maximal ‘push’,^38^). A value close to 0 indicates low bias whereas values greater than 1 mean that the bias is greater than the user’s maximum effort. We measured normalized bias across all pairs of sessions by training an offline linear decoder on a reference session and then measuring its normalized bias index on a test session (Fig. 1E). We also include bias estimates within-session when using decoders on both training and holdout blocks, which had average normalized bias values of 0.117 0.154 and 0.241-0.257 (respectively, mean±SD). After 1-2 weeks, median bias values rose to around 0.79, which is high enough to render control almost impossible.

We then evaluated PD changes by examining the cosine angle between linear decoder weights on different pairs of sessions (“subspace drift”). This metric provides a global measure of all PD changes occurring across the neural population (see Methods), with a value close to 1 when readout dimensions are highly aligned and close to 0 when orthogonal. We then plotted the cosine angle as a function of the gap between sessions (Fig. 1F).The median tuning similarity within-session was around 0.75 (43 degree difference) but dropped with 1-2 weeks of separation between sessions (71 degree difference). The median R^2^ correspondingly moved from 0.39 to 0.21, highlighting the impact of nonstationarities on downstream task performance. These results highlight how nonstationarities can quickly accumulate and impact decoder performance. While mean shifts can easily be accounted for (e.g with adaptive offset-correction methods as in ^5^), PD changes present a more advanced challenge to long-term robustness that motivated us to design PRI-T.

### Probabilistic target inference via a hidden Markov model

Strategies such as FA stabilization (Fig. 2A) and ADAN aim to stabilize decoders in the face of PD changes by taking advantage of a subset of consistent neural activity patterns that may persist across days. In the case of FA stabilization, this approach involves training a FA subspace decoder on some reference day, and then using a Procrustes alignment to realign the FA model from a new day (Fig. 2B).

**Figure 2.**
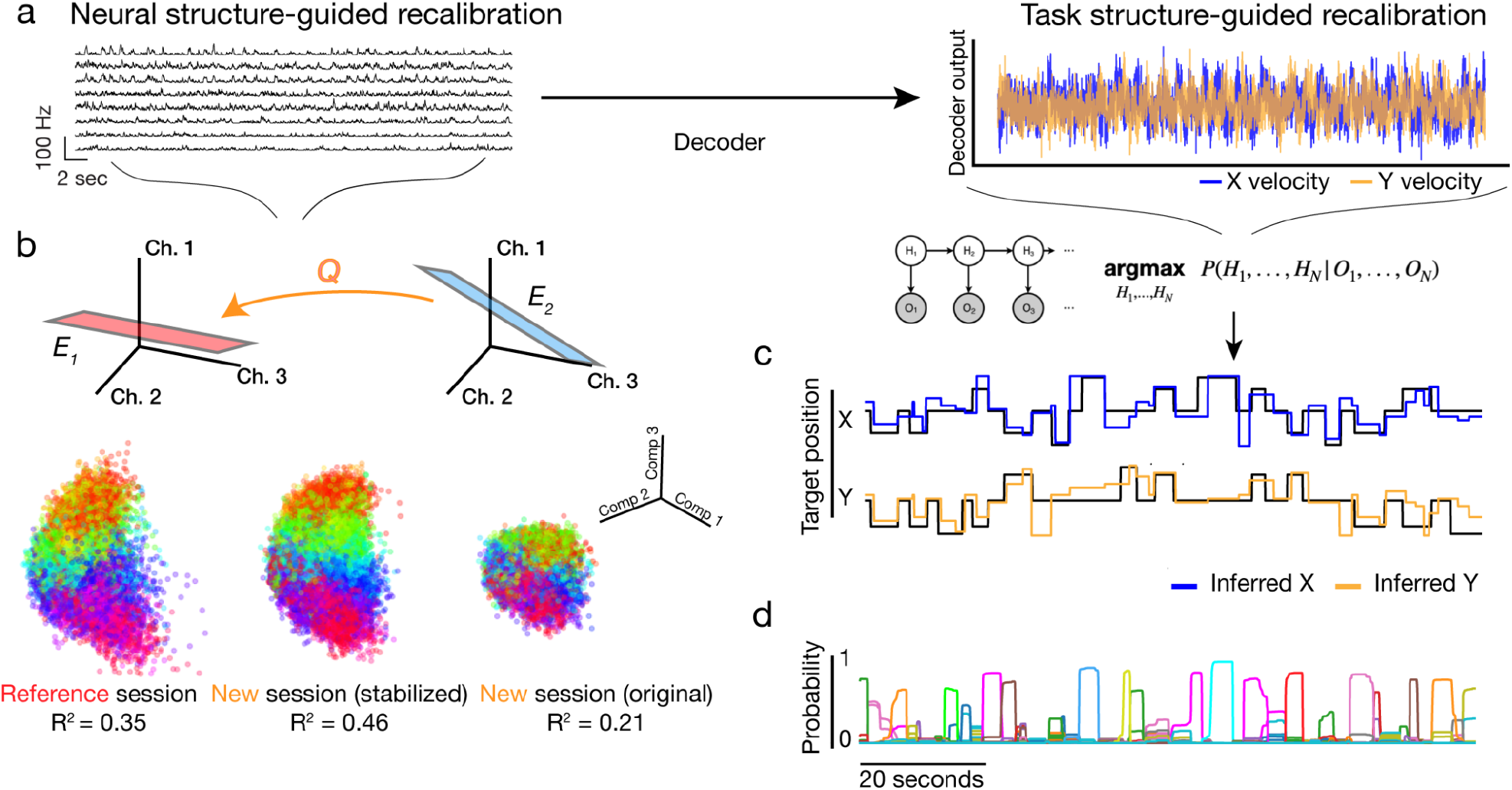
FA stabilization and PRI-T overviews. **(a)** Neural firing rates on a new day are fed through an old decoder, resulting in a noisy cursor velocity command timeseries. **(b)** FA stabilization identifies an orthogonal map Q that realigns a new day’s subspace E’ with the reference day, E. State-space plots show neural activity in top FA components, colored by cursor displacement angle. A decoder trained in the reference latent space provides decent performance, but drops in performance when fed data from a new session (“original”). Applying stabilization via Q increases performance by aligning the subspaces (“stabilized”). **(c)** PRI-T instead operates on decoder outputs, by identifying the most likely (Viterbi) sequence of targets {*H_t_*} given the observed velocities and cursor position {*O_t_*}, which closely matches the ground-truth. **(d)** We also query the model for its confidence in its predictions by obtaining the marginal probability of the target state *P*(*H_t_*|*O_1_*,…,*O_N_*) and use this to weight the importance of each pseudo-target label during recalibration.

Alternatively, one might use task-related structure to perform unsupervised recalibration (Fig. 2A, right) by inferring the user’s intended targets from noisy decoder outputs, and then using those inferred targets to retrain the decoder. To optimize this process, we would ideally like a labeling model which provides both 1) pseudo-labels for retraining our cursor decoder and 2) an associated measure of confidence in its outputs.

RTI is one task structure-based approach, and assumes that user selections happen when the cursor is hovering over a target ^5^. Data from a preceding time window, when the user is presumably moving toward estimated target locations, is used to update the decoder after the target selection occurs. RTI is hence unsupervised but its pseudo-labels don’t have associated confidence estimates, which might be helpful for weighting the importance of a given target estimate to the overall recalibration. Further, there is additional information in the cursor trajectory that can’t be leveraged by examining the user’s intended clicks alone. For example, If one observes a cursor moving steadily toward a given corner of the screen we might infer that a user is aiming toward a target there. This provides complementary information that can be exploited for better target labeling. RTI also requires the user to be actively selecting targets to enable retrospective retraining, rendering the approach unsuitable for situations where a user might passively hover over parts of the screen (e.g. drop-down menus, browsing).

We leverage continuous cursor information, and optionally discrete selections, by linking them to the target state via PRI-T, a probabilistic model (see Methods). PRI-T uses a Hidden Markov Model (HMM) to obtain target estimates in a principled way, enabling continuous predictions alongside confidence estimates (Fig. 1C) that allow us to differentially weight the importance of different target estimates for recalibration (Methods). Using a probabilistic model unlocks a wider array of downstream algorithms (e.g. weighted least squares) and easy integration of auxiliary signals such as click when available.

### Sensitivity of existing recalibration strategies to long-term PD changes

To benchmark PRI-T against existing approaches, we examined robustness across multiple timescales via offline analysis of session pairs. Since neural nonstationarities generally increase with larger gaps between sessions (Fig. 1E,F), it is possible that recalibration systems might perform well in the short-term, when such changes are small, but fail in the long-term.

We analyzed this possibility using a cursor control dataset from participant T5 spanning roughly five years and generated pairs of sessions (see Methods) for evaluating recalibration performance. For each pair, a decoder was trained on the earlier session and tested on the newer session after separately applying five methods: supervised recalibration, mean recalibration, FA stabilization, ADAN, and PRI-T. Plotting performance across pairs of sessions as a heatmap (Fig. 3A), we observed similar overall performance patterns across approaches. For a more direct comparison, we normalized scores against supervised recalibration’s performance and plotted the data side-by-side (Fig. 3B). All methods generally outperformed mean recalibration (0.78, [0.70-0.87]; median, IQR), implying that all methods were successfully accounting for some of the PD changes as well as mean changes on this subset of days. FA stabilization, ADAN, and PRI-T all clustered around 0.89-0.91 in median scores but with variable spread. Both latent space methods tended to have much larger IQRs ([0.83 - 0.96] for stabilizer; [0.89 - 0.98] for ADAN) compared to PRI-T ([0.85 - 0.96]). We also tested a simple ensembling model, which averages decoder predictions from PRI-T and FA stabilization, that provided the highest relative performance across methods (0.96, [0.93 - 0.99] median and IQR; Fig. 3B) in this single recalibration setting. Pairwise comparisons across methods had similar findings, with all unsupervised methods outperforming mean recalibration and otherwise appearing equivalent aside for the ensembled approach (Methods; Fig. S3). These results were consistent when examining R^2^ as well (Fig. S4).

**Figure 3.**
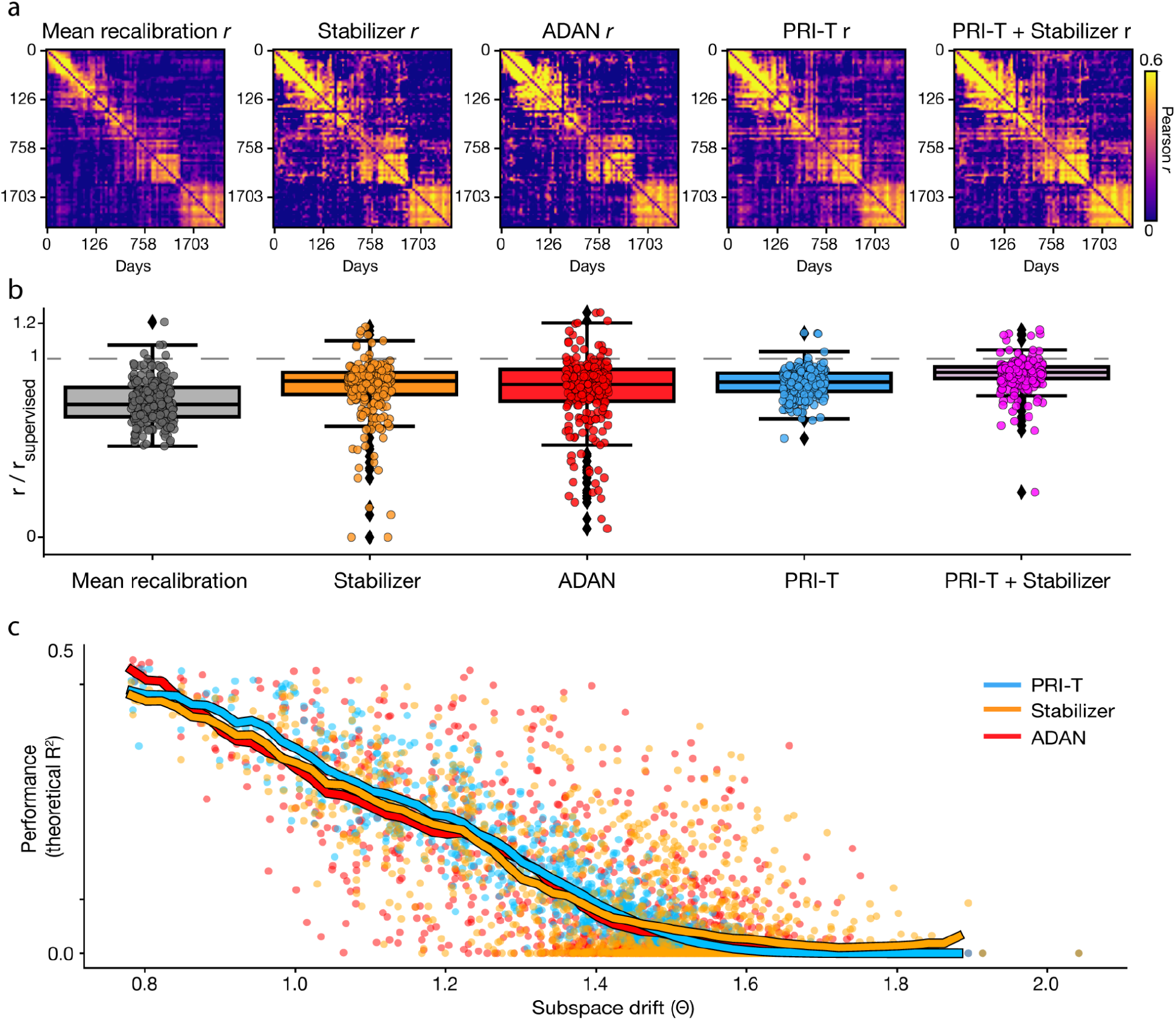
All tested methods fail with sufficient tuning drift between sessions. **(a)** Performance of various approaches across different pairs of sessions and plotted as a heatmap. Color indicates decoder output correlations with ground-truth displacement after recalibration. **(b)** Relative performance compared to supervised recalibration (1 = equivalent to supervised). **(c)** Performance (r^2^, see Methods) as a function of the angle between supervised decoder weights on reference and new days. Solid line corresponds to moving average.

Interestingly, all approaches had similar failures, underperforming on day pairs that are further apart in time (upper right corners in Fig. 3A) compared to when sessions were closer to one another (banding pattern along the diagonal in. Fig. 3A). Since far-apart sessions likely have a much larger degree of signal nonstationarity compared to closer sessions, we reasoned that this trend may reflect an inability to compensate for long-term signal drift. We explicitly tested this hypothesis by analyzing performance as a function of subspace drift. Results show that PRI-T, ADAN, and FA stabilization all perform poorly in the face of large rotations of the subspace in which neural tuning to movement exists, and seem to fail completely for rotations larger than roughly 90 degrees (albeit with differing variability; Fig. 3C).

These initial offline findings indicate that, when performing unsupervised recalibration using only a single pair of days, all approaches appear unsuitable for maintaining long-term cursor control in our participant. On a shorter timescale, however, performance was somewhat more stable. We therefore reasoned that iterative recalibration approaches that use chains of days might work better to counter nonstationary neural activity. Our intuition here was that chopping up long-term neural changes into a series of smaller day-to-day changes would ultimately be more tractable, owing to the less variable differences between nearby days.

### Building a realistic closed-loop model of neural drift

To examine the hypothesis that daily, unsupervised recalibration fares better against long-term signal changes, we would like to obtain long stretches of consistent cursor control data without major time gaps. Since we are also interested in evaluating various recalibration strategies, we require a tractable approach that does not require too much valuable session time with clinical trial participants.

Consequently, we turned to a closed-loop simulation environment to enable large-scale, systematic performance comparisons across many recalibration strategies. We used a variant of the PLM simulator from ^38^ with an additional neural tuning model and matched SNR characteristics to T5’s neural data (Fig. 4A-B, see Methods). We also matched the overall course of subspace drift, defined as changes in the encoding subspace orientation across time, by fitting an exponential decay model to encoding subspace drift over two weeks for the X and Y-subspaces separately (Fig. 4C, left panel). This yielded decay constants of 0.91/day and 0.93/day for the X and Y-subspaces respectively. As these values are relatively close, we opted to use the lower of the two values for both X and Y-subspace drift (Fig. 4C right).

**Figure 4.**
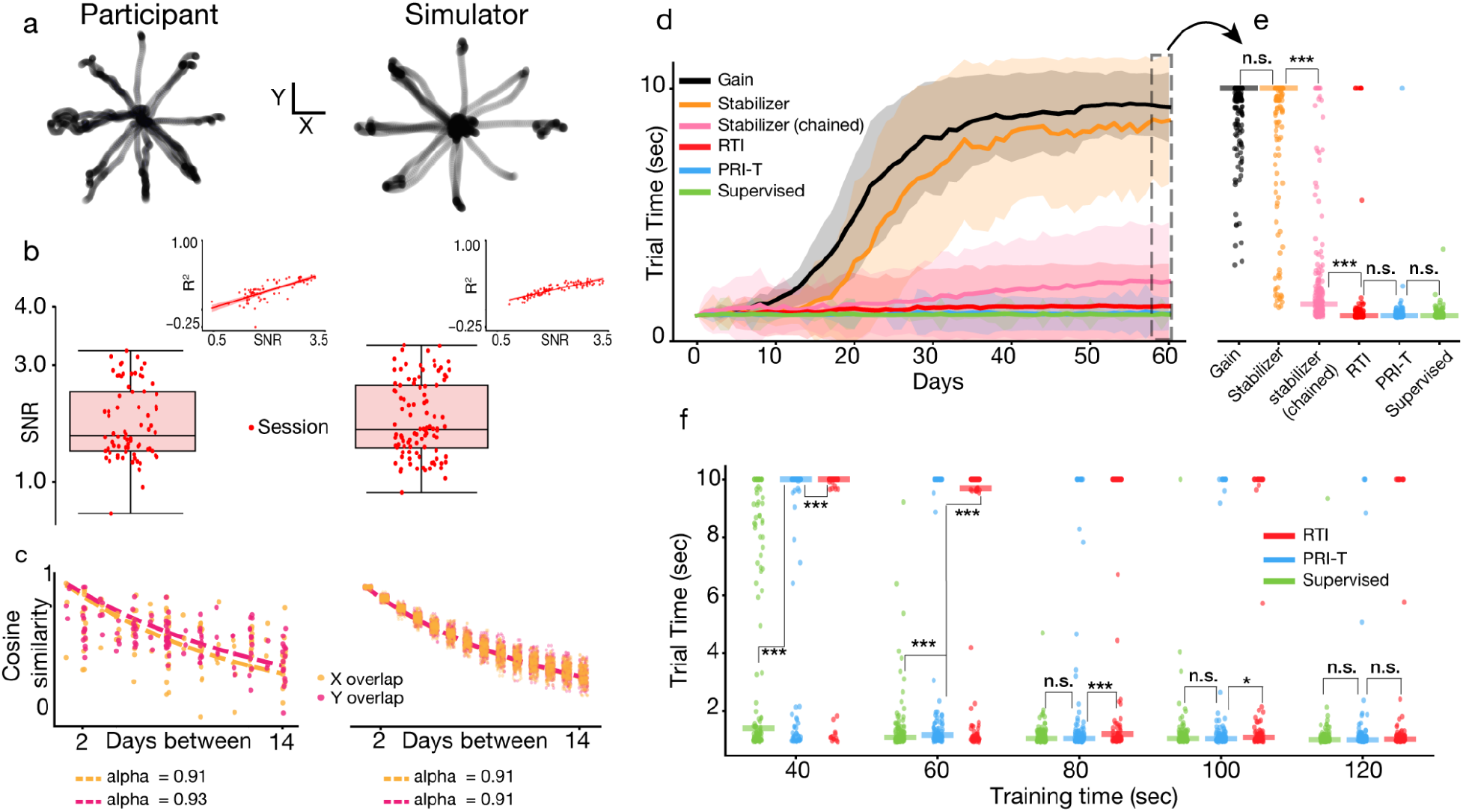
Comparison of recalibration strategies in simulation. **(a)** Example trajectories during real (left) and simulated (right) closed-loop BCI use. **(b)** Distribution of SNR values in participant T5 (left) and simulation (right). T5’s SNR distribution (median 1.85, 1.53-2.59 IQR) closely matches the simulator (median 1.97, 1.53-2.65 IQR). For simple visual comparison, we plot 100 datapoints drawn from the simulated SNR distribution. Insets show correlation with offline R^2 during heldout timepoints (shading: 95% confidence intervals). Median R^2^s were close between the two settings (0.28 vs. 0.30 for experimental and simulated) and with similar IQR ranges (0.18 - 0.40 experimental, 0.22 - 0.37 simulated). **(c)** Cosine angle difference of tuning models across separate sessions in T5 (left) and simulation (right). Dotted lines show exponential decay fits. **(d)** Average trial times across sessions for different methods (400 seconds or 6.7 minutes of data per block). Shading corresponds to standard deviation over 200 independent runs. (**e**) Comparison of trial time values between methods at 60 days out. **(f)** Performance of RTI and PRI-T as a function of training data size with 100 independent runs each per data size tested. Trial times measured at 30 days out.

This latter step ensures similar average drift between simulated and observed neural activity, but leaves unspecified the variance of this drift. We chose to simulate a low variance in the amount of day-to-day drift, motivated by progress in next-generation BCI systems ^39–41^. As channel counts scale, iBCIs will likely acquire a degree of signal redundancy that provides some protection from nonstationarities (as noted in e.g. ^42^). More plainly, this idea captures the intuition that, for example, a decoder built from 5 channels will have more variable performance when 2 channels shift than a cursor decoder built from 500 channels where 200 shift. Each channel’s drift perturbs the overall encoding subspace, but their combined effect tends to wash out extreme scenarios ^9,42^. This implies that high-channel count BCI systems should exist in a regime where consistent drift occurs across time but with limited variability, making the case for adaptive algorithms that can compensate for these changes. We demonstrate this in a toy model with a simple linear encoding system (see Methods), where we show empirically and with a rough proof that variance falls off with increasing channel counts (Fig. S6A-C).

### Simulations show that only target labeling strategies maintain long-term control

We then simulated closed-loop control over two months in the presence of a drifting encoding subspace. For all decoders, we optimized the cursor gain within each day so that we could more closely measure performance related to how well the decoder tracks this subspace (versus, for example, variability arising from high-speed cursors overshooting targets; see Methods). As changes compound, a decoder fit on the first day (black curve, Fig. 4D) grows increasingly noisy until it is no longer usable (average trial time = 9.28 1.39 seconds/trial at 60 days out, mean ± SD). This breakdown happens despite gain optimization within each day, implying that poor performance is due to low-quality decoder weights. By contrast, using supervised recalibration (green curve) yields higher and more consistent performance over time with an average trial time of 1.05 ± 0.22 at 60 days (p < 1e-60, Wilcoxon rank-sum test).

We also simulated three unsupervised recalibration procedures: PRI-T, RTI, and FA stabilization (with hyperparameter searches for all, see Fig. S1). We found that FA stabilization was prone to noise as time passed between the reference and new day’s data (black curve), eventually yielding median performance similar to gain optimization alone, with trial times of 8.73±2.52 seconds (p = 0.57, Wilcoxon rank-sum).

To avoid this issue, we reasoned that on a short timescale, drift is minimal and sufficiently many stable channels should exist to make the alignment problem more tractable. We hence implemented a “daisy-chained” FA stabilizer, which leverages relatively easy day-to-day mappings to stabilize neural data with respect to the reference day (pink curve); rather than mapping a new session’s activity to the original subspace, we instead learn a set of easier day-to-day maps and compose them to obtain a transformation into the original subspace (Fig. S1E). This approach proved far more robust over long time spans compared to the standard stabilizer (p < 1e-50; Wilcoxon rank-sum) but yielded suboptimal control, with some sessions being totally uncontrollable (session dots with average trial time values at ~10 seconds, Fig. 4D) and a 2x slower average trial time than supervised retrained decoders.

By contrast, PRI-T and RTI enabled consistently stable control and outperformed daisy-chained FA stabilization (p < 1e-34 both, Wilcoxon-rank sum test). Both approaches had similar performance (1.37±1.68 RTI, 1.09±0.65 PRI-T) and, encouragingly, did not differ significantly from supervised recalibration at the end of the simulated stretch (p > 0.18 both, Wilcoxon-rank sum test). Hence PRI-T and RTI were the only two strategies that enabled high-quality, long-term control for the simulated user.

### *In-silico* data efficiency of PRI-T and RTI

As RTI and PRI-T were close in performance, we then looked at the data efficiency of each method by measuring performance after 30 days of unsupervised recalibration while varying the amount of training data provided. PRI-T was far more robust to smaller dataset sizes, with significantly faster trial times across most sizes compared to RTI (p < 0.05 all for 40-120 seconds of training data, Wilcoxon rank-sum test; Fig. 4F) and similar performance to supervised recalibration (p > 0.05 all except 40 and 60 seconds); PRI-T maintained qualitatively similar control to supervised recalibration at even a 60% reduction in the amount of training data (80 seconds training size, p = 0.52, Wilcoxon rank-sum test).

We also evaluated the data efficiency of click-augmented PRI-T (see Methods), where an explicit click observation model is integrated into the HMM. This resulted in a significant improvement compared to the standard PRI-T model at smaller data size (p < 0.05 for 40, 60, and 80 second sizes; Fig. S4A), validating the integration of click and cursor decoder information to improve performance beyond either signal alone.

### PRI-T demonstrates superior performance during one month of closed-loop control with T5

PRI-T’s superior performance in simulation was encouraging, but nevertheless may not necessarily generalize to real closed-loop control in T5. We therefore tested the ability of PRI-T to enable fully unsupervised, closed-loop control as T5 engaged in a random target task over one month.

During each session, the weights of a linear decoder were updated using PRI-T after each closed-loop block (see Methods for decoder details). We benchmarked PRI-T’s ability to compensate for neural changes by also comparing FA stabilization and a fixed decoder with bias subtraction to account for mean shifts (Methods). PRI-T consistently outperformed the other decoders, both near the onset of closed-loop data collection and toward the end. PRI-T’s advantage increased considerably when tuning diverged the most from the reference day. By contrast, FA stabilization was highly variable and often failed to improve upon a fixed decoder (e.g. day 12). However, FA stabilization outperformed the fixed decoder on the last day, when neural activity presumably reverted back to a similar pattern as on day 0, evidenced by the improved performance of the fixed decoder between days 14 (6.83 ± 3.03 seconds per trial; mean ± SD) and 28 (3.61±2.12).

We also collected an open-loop (OL) block at the beginning of each day to better probe decoder performance without closed-loop effects (Fig. 5C). All trends were roughly consistent, with PRI-T outperforming FA stabilization further out in time (days 7, 12, and 14). On day 28, PRI-T was similar to FA stabilization while both methods outperformed the fixed decoder. The fixed decoder partially recovers on day 28, implying that the neural activity dimensions found on day 28 have stronger overlap with day 0 than the prior sessions. This is possible given that neural activity evolves in a somewhat random walk fashion, and may thus double back on itself. Nevertheless, PRI-T appears to be the best method for adapting to nonstationarities, as a majority of the time, neural activity will differ substantially from an initial state as time progresses.

**Figure 5.**
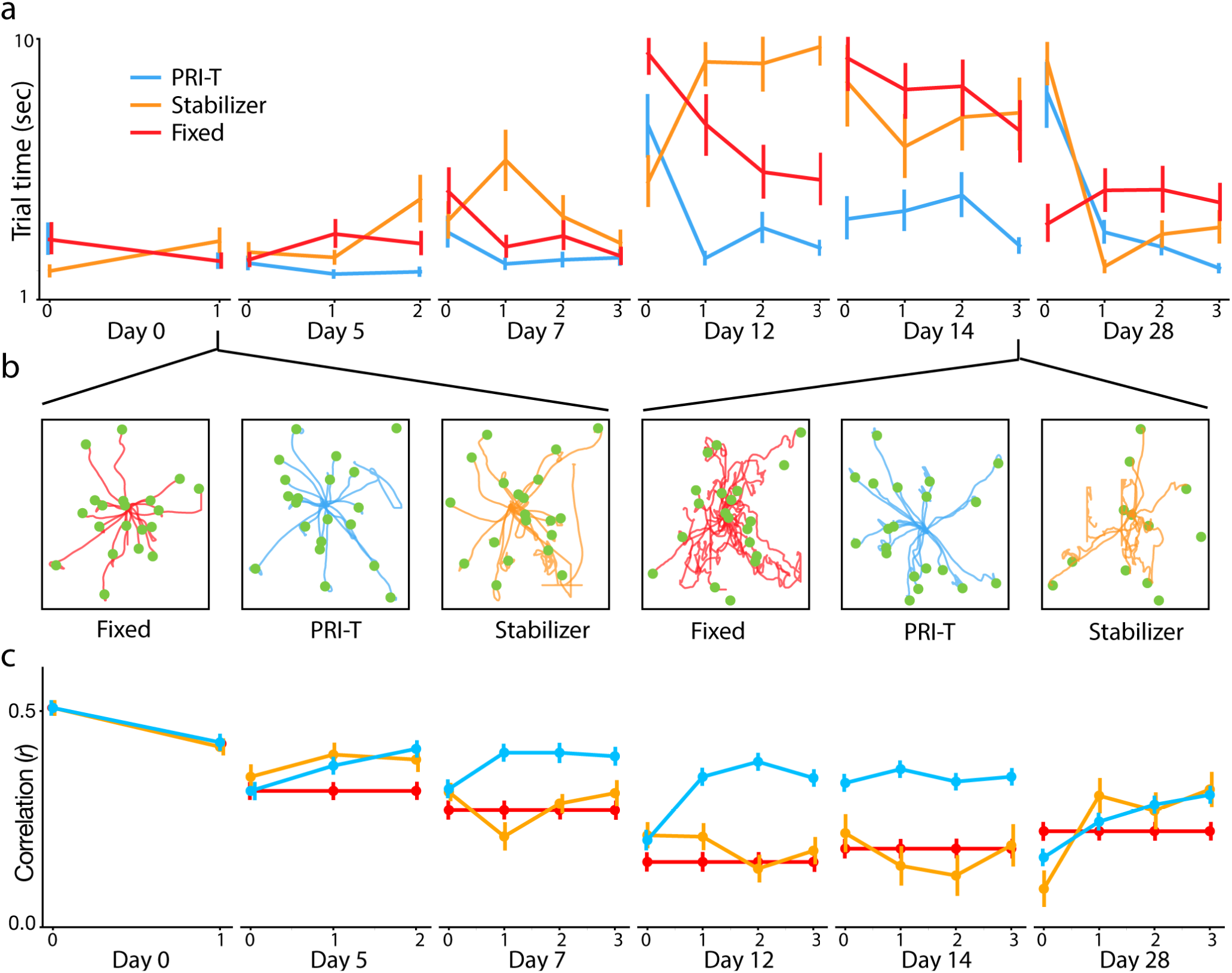
Closed-loop performance comparison. **(a)** Average trial times across blocks and sessions for three different decoder strategies (bootstrapped empirical 95% confidence intervals). After initial supervised training on day 0’ the decoders were tested and recalibrated repeatedly in an interleaved manner throughout a session. Ticks represent rounds of testing and track the number of prior recalibrations on a given day (e.g. day 7, slot 3 tracks performance across decoders after 3 prior iterations of deployment and recalibration). On a new day, the decoders from the end of the prior day are loaded for round 0. (**b**) Example cursor trajectories from the first and fifth sessions for all three recalibration strategies. Green dots are target locations. Trial starts are recentered to the origin. (**c**) OL block correlations for each decoder. Intervals again reflect 95% confidence intervals (empirical bootstrap). Throughout the second week, PRI-T outperformed FA stabilization in OL decoding (an average 0.14 0.078 increase in correlation, or 84% relative improvement on average).

## Discussion

Our offline, *in-silico*, and online results demonstrate an unsupervised recalibration system that outperforms prior state-of-the-art and helps tackle one of the remaining technical roadblocks for clinical translation of cursor BCI systems. By providing consistent control without supervision - despite compounding neural nonstationarities - these findings demonstrate a way for BCI users to maintain robust control without frequent pauses for supervised recalibration. Our work also demonstrates that iterative or “chaining” based strategies may be necessary to compensate for long-term signal drift and that, at least in simulation, only target-inference methods appear capable of indefinite chaining.

Recent work has examined neural latent space recalibration procedures ^6,25,27,28^. In studies with non-human primates (NHPs), these approaches align neural activity to a reference day, enabling a fixed decoder to continue performing despite accruing neural changes. However, In our offline analysis, we found that two such approaches - FA stabilization and ADAN - were unable to consistently realign activity in the face of large subspace drifts (Fig. 3C). ADAN had similar performance to FA stabilization although with larger variability, which may reflect a tendency to overfit on these data (Methods). Similarly, PRI-T was unable to restore performance when subspace changes were large between reference and test sessions. We suspect these failures reflect an inability to deal with long-term drift, where the encoding subspace has moved far away from an initial reference location. In the case of FA stabilization, the alignment step may fail if the underlying task manifold is too symmetrical to enforce a unique identification (a failure mode seen in other subspace alignment methods e.g.^23,29^ with kinematic and neural distribution realignment, respectively). For instance, linear tuning to velocity-like signals can cause a radially symmetric distribution of neural activity within the task subspace. If few stable channels exist to constrain the Procrustes problem, then the alignment may be particularly sensitive to rotations. This aligns with recent work showing that both FA stabilization and ADAN underperformed with NHP center-out reaching data ^28^. More immediately, this failure mode would explain FA stabilization’s poor performance offline on pairs of sessions with large subspace angle changes (Fig. 3), in our closed-loop simulations (Fig. 4), and during closed-loop control in T5 on day 12, which had the largest neural nonstationarities with respect to day 0 (Fig. 5).

Since long-term drift rendered all approaches ineffective, we next reasoned that iterative updates are required to maintain high performance. We demonstrate this in simulation using a variant of FA stabilization (“chained”, Fig. 4), where subspace mappings are fit across consecutive pairs of sessions and then chained together. Despite stability improvements, chained FA stabilization still showed overall high variability, with only target inference approaches (RTI and PRI-T) enabling consistent long-term control in simulation. In real closed-loop control, applying PRI-T iteratively also enabled relatively stable performance across one month. Chaining FA stabilizers in simulation likely failed due to compounding errors over time. If each iteration of the stabilizer introduces an independent estimate of the optimal rotation angle from day-to-day, then the final transformation (which is composed of these smaller mappings) will accumulate errors over time without a self-correction mechanism.

Instability of neural signals may be greater in humans than in animal models such as NHPs. For instance, one offline NHP decoding analysis demonstrated consistent performance over hundreds of days using fixed linear decoders (Chestek et al. 2011). Yet our work shows little correlation on average between task subspaces after a few months (Fig. 1) and highly variable performance in closed loop with FA stabilization. Given the single subject nature of the study, one hypothesis for the discrepancy is that T5’s neural signals are unusually unstable relative to other intracortical BCI users. However, other work has identified rapid fall-offs in the number of stable channels across time ((Downey et al. 2018)), albeit with some highly stable subsets, as well as closed-loop instability in human users ^3,5^.

While recent work has often focused on neural distribution-matching strategies, many of these have yet to be evaluated for online BCI control. One exception is FA stabilization, which was evaluated over a 5 day period in NHPs ^6^, and may be optimal for a specific noise regime experienced in shorter timeframes. By contrast, self-training has shown much promise with Li and colleagues ^26^ demonstrating robust cursor control over 29 days in NHPs using bayesian updates on decoder weights. Similarly, Jarosiewicz and colleagues ^32^ deployed RTI with a human iBCI participant for stable cursor control over 30 days. Alongside our closed-loop results here (28 days), we believe that self-training strategies demonstrate strong potential for long-term BCI stability.

Strategies such as PRI-T may therefore be crucial for long-term stability in human recordings. Future efforts to quantify the extent of signal nonstationarity differences between animal models and human BCI users may enable a more nuanced understanding of the conditions in which recalibration procedures will generalize to human users. By leveraging relatively simple task structure, unsupervised retraining procedures can simplify and stabilize long-term control for end users, aiding clinical translation of cursor BCIs.

## Author Contributions

We have included an author contributions matrix graphic inspired by the recent, forward-looking piece in Nature Index ^43^.

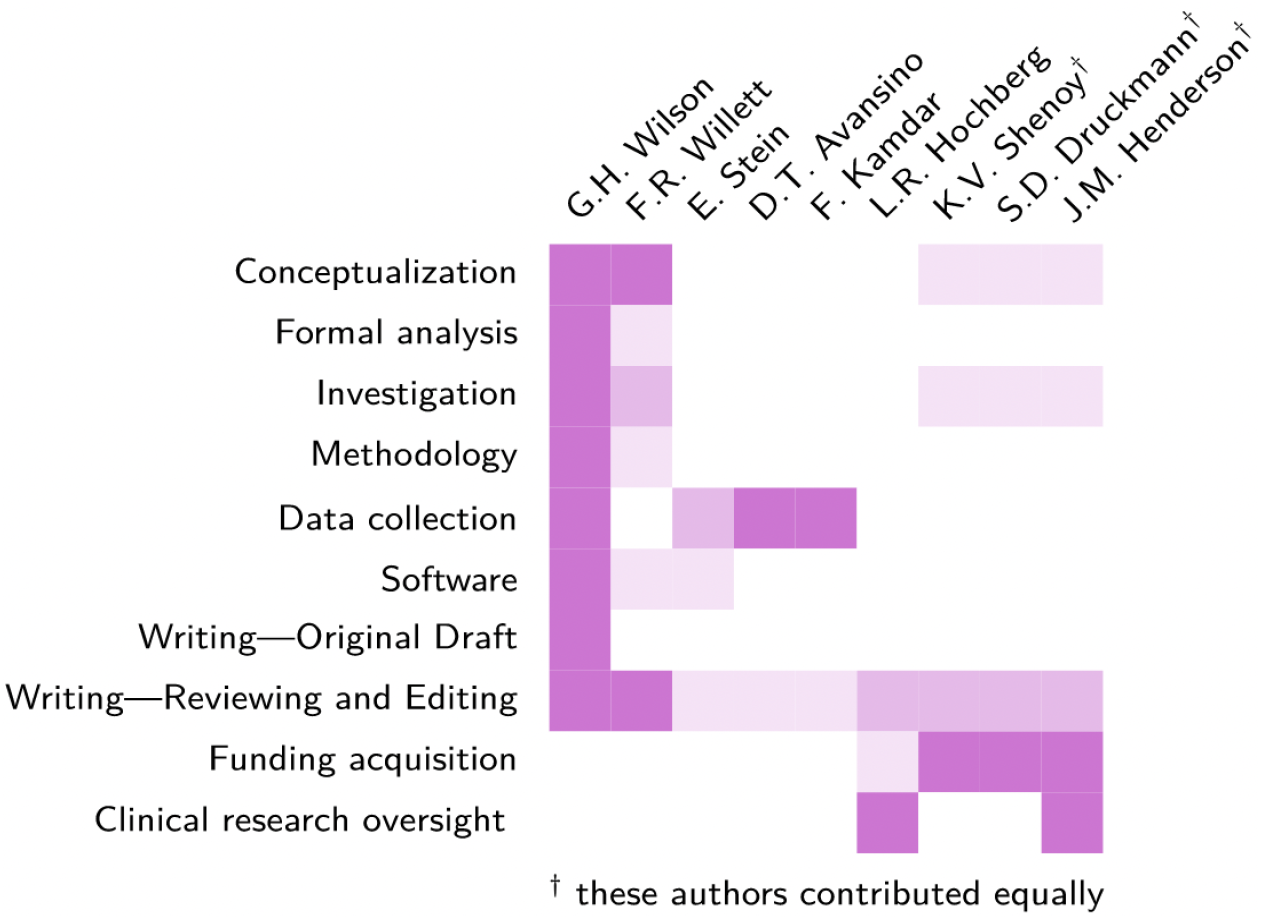

## Acknowledgements

We thank participant T5 and his caregivers for their generously volunteered time and effort as part of our BrainGate2 pilot clinical trial. We also thank our Stanford Neural Prosthetics Translational Lab, Neural Prosthetic Systems Lab, and the Druckmann lab groups for helpful discussions. We thank Beverly Davis, Sandrin Kosasih, and Kathy Tsou for expert faculty-administrative and lab management support. Some of the computing for this project was performed on the Stanford High Performance Computing Cluster (Sherlock cluster). We thank Stanford University and the Stanford Research Computing Center for providing expert IT support for this computational resource where our primary compute nodes are housed and maintained.

This work was supported by an NSF Graduate Research Fellowship DGE-1656518 and Regina Casper Stanford Graduate Fellowship (G.H.W.); Howard Hughes Medical Institute (HHMI) at Stanford (F.R.W., D.T.A); NIH-NIDCD U01DC017844, NIH-NINDS U01-NS098968, Larry and Pamela Garlick, Wu Tsai Neurosciences Institute at Stanford (J.M.H and K.V.S); Department of Veterans Affairs Rehabilitation Research and Development Service A2295R and B6453R (L.R.H); NIH-NIDCD R01DC014034 (L.R.H, J.M.H, K.V.S); NIBIB R01-EB028171, Simons Foundation Collaboration on the Global Brain 542969 (S.D.); and Simons Foundation Collaboration on the Global Brain 543045, Hong Seh and Vivian W. M. Lim endowed chair and HHMI Investigatorship at Stanford University (K.V.S).

---

**Supplementary figure 1.**
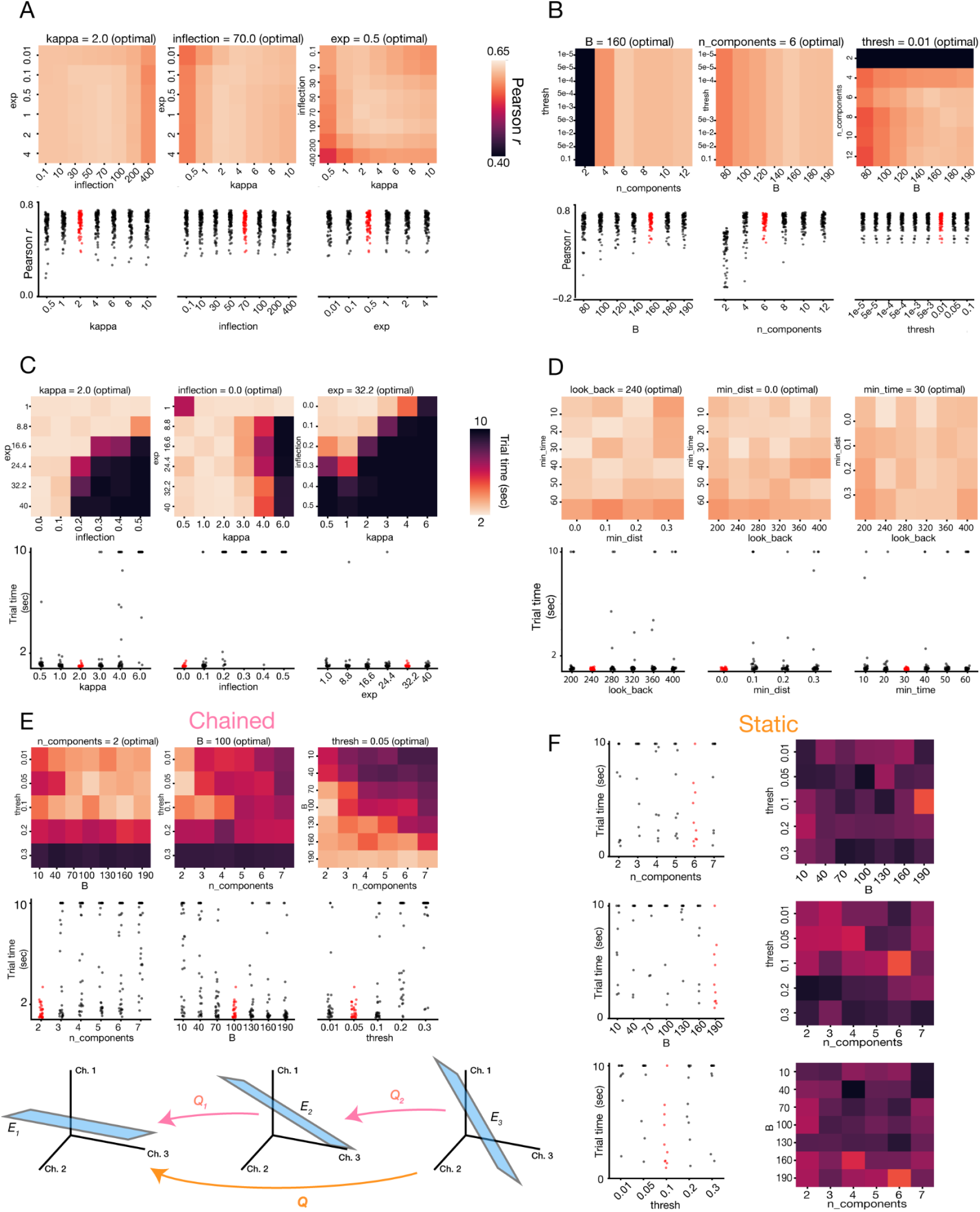
Hyperparameter sweeps. **(A)** Offline PRI-T hyperparameter (HP) sweeps. Top: 2D heatmaps of median Pearson correlations as a function of HPs (left-out value is set to optimal). Bottom: Correlation values while varying one HP at a time (others are set to their optimal values). Highlighted red scatters indicate optimal values in sweep (maximizing median correlation). **(B)** Offline subspace stabilizer sweeps. (**C-D**) Simulator HP sweeps: all sweeps done using 30 independent runs for each HP set and evaluated at 30 days with average trial time. (**C**) Simulator PRI-T HP sweeps. (**D**) Simulator RTI HP sweeps. (**E**) Simulator FA stabilization sweeps, where stabilizer is daisy-chained. (**F**) Simulator stabilizer sweeps using standard (“static”) approach.

**Supplementary figure 2.**
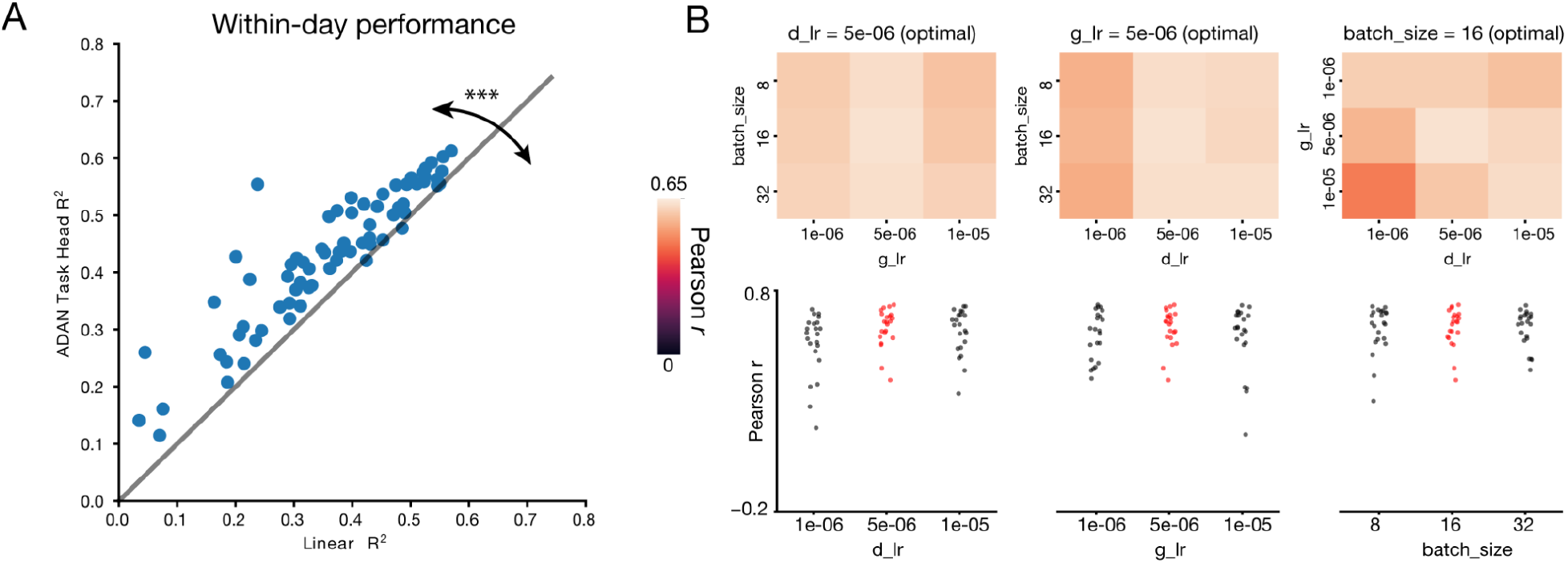
ADAN details. **(A)** ADAN models improved on linear regression by 23% on average within-day (p < 0.001, Wilcoxon signed-rank test). **(B)** Offline ADAN hyperparameter sweeps. Top: 2D heatmaps of median Pearson correlations as a function of hyperparameters (left-out value is set to optimal). Bottom: Correlation values while varying one hyperparameter at a time (others are set to their optimal values). Highlighted red scatters indicate optimal values in sweep (maximizing median correlation).

**Supplementary figure 3.**
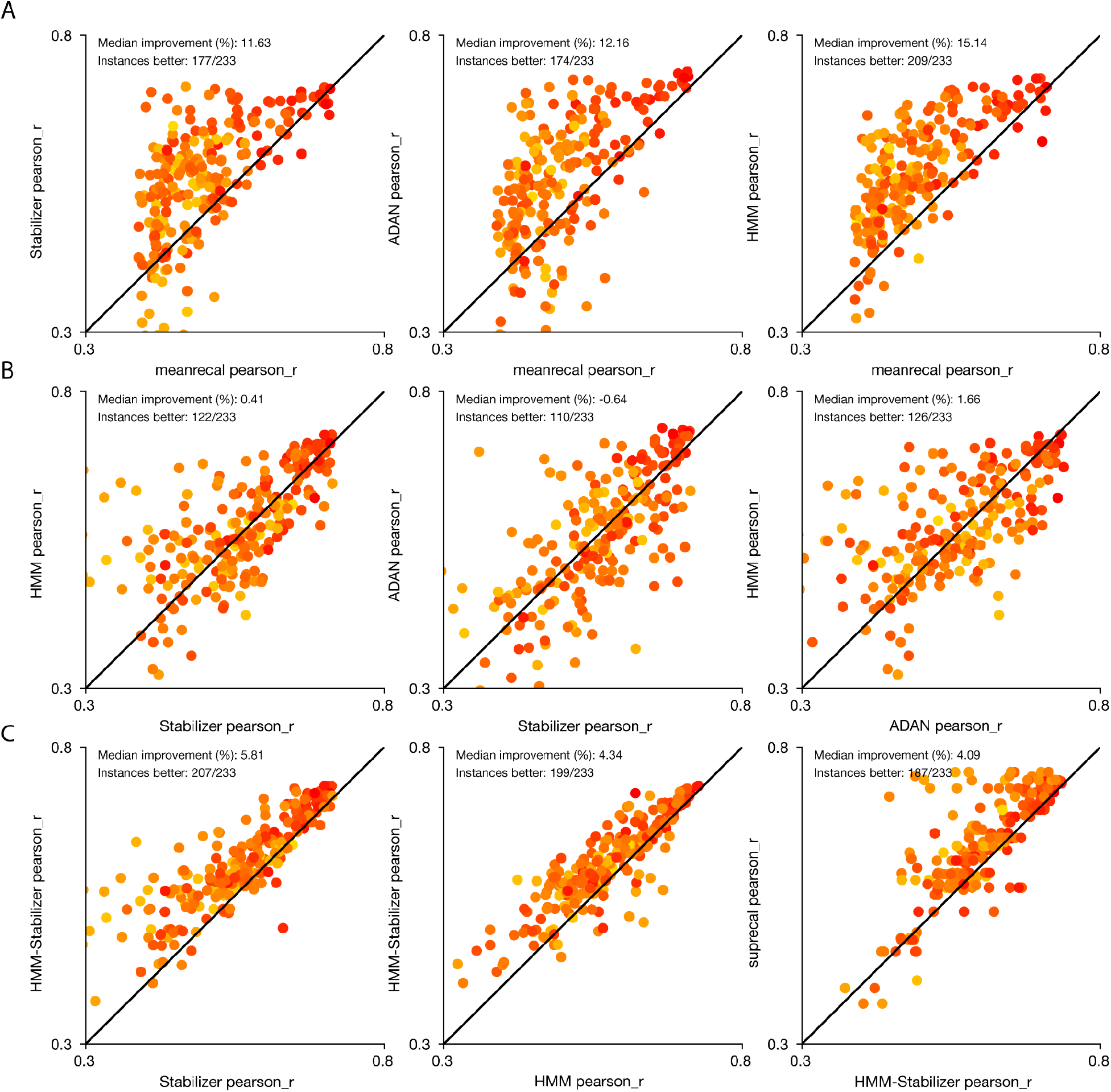
Pairwise offline results (Pearson correlation). **(A)** Pairwise comparisons plotted for PRI-T, FA stabilizer, ADAN against mean recalibration. PRI-T, stabilizer, and ADAN all generally outperform mean recalibration (> 75% of cases for all three). (**B**) Comparisons among PRI-T, ADAN, and FA stabilizer. Methods are all roughly equivalent. (**C**) Combined approach outperforms versus PRI-T and stabilizer individually (>85% of the time in both cases). Combined decoder is slightly worse than supervised recalibration (supervised outperforms in ~80% of cases with a median 4% improvement).

**Supplementary figure 4.**
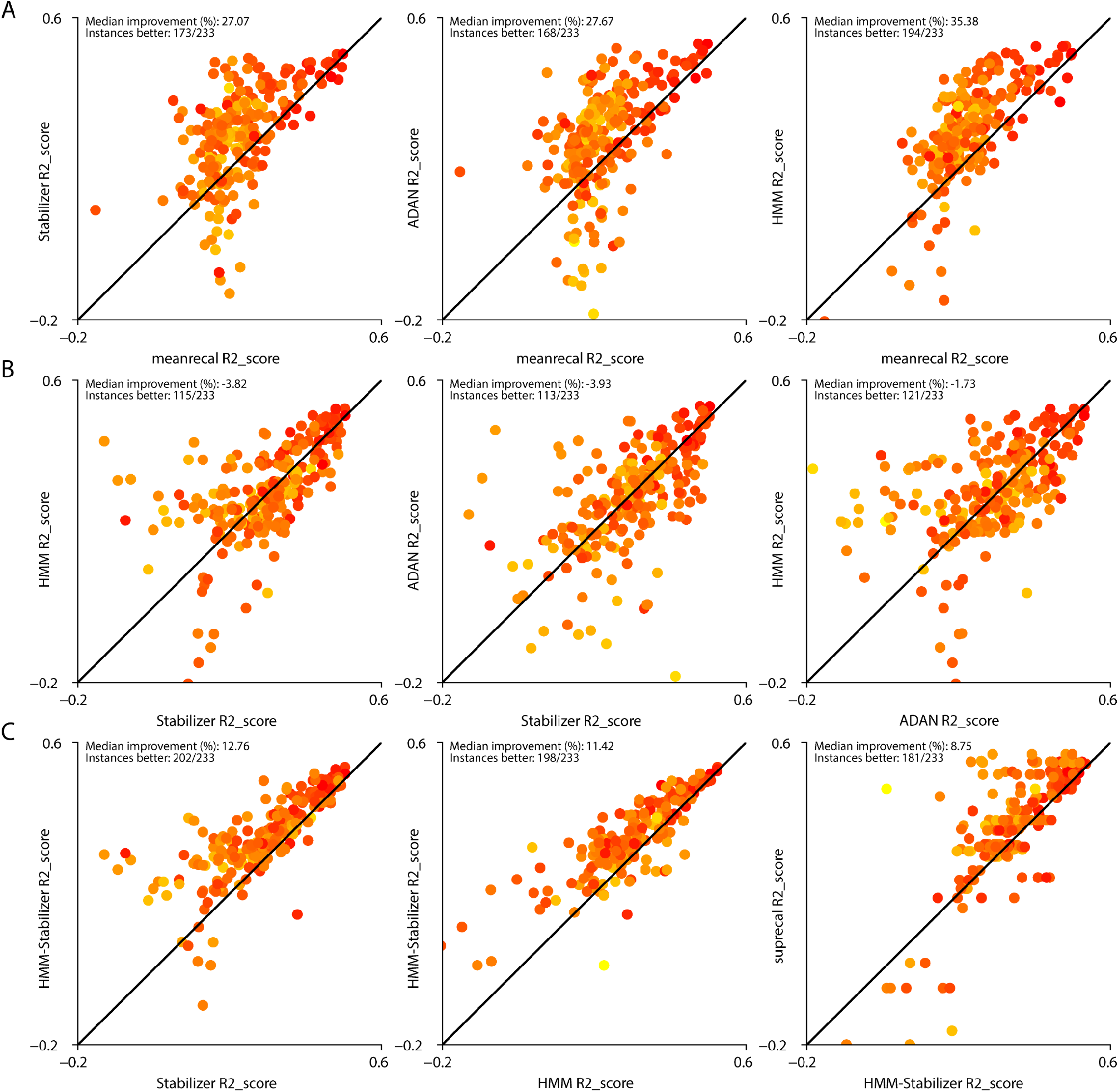
Pairwise offline results (R^2^). Same as S3 but with R^2^ score. **(A)** Pairwise comparisons plotted for PRI-T, FA stabilizer, ADAN against mean recalibration. PRI-T, stabilizer, and ADAN all generally outperform mean recalibration (> 72% of cases for all three). (**B**) Comparisons among PRI-T, ADAN, and FA stabilizer. Methods are all roughly equivalent. (**C**) Combined approach outperforms versus PRI-T and stabilizer individually (>84% of the time in both cases). Combined decoder is worse than supervised recalibration (supervised outperforms in ~77% of cases with a median 9% improvement).

**Supplementary figure 5.**
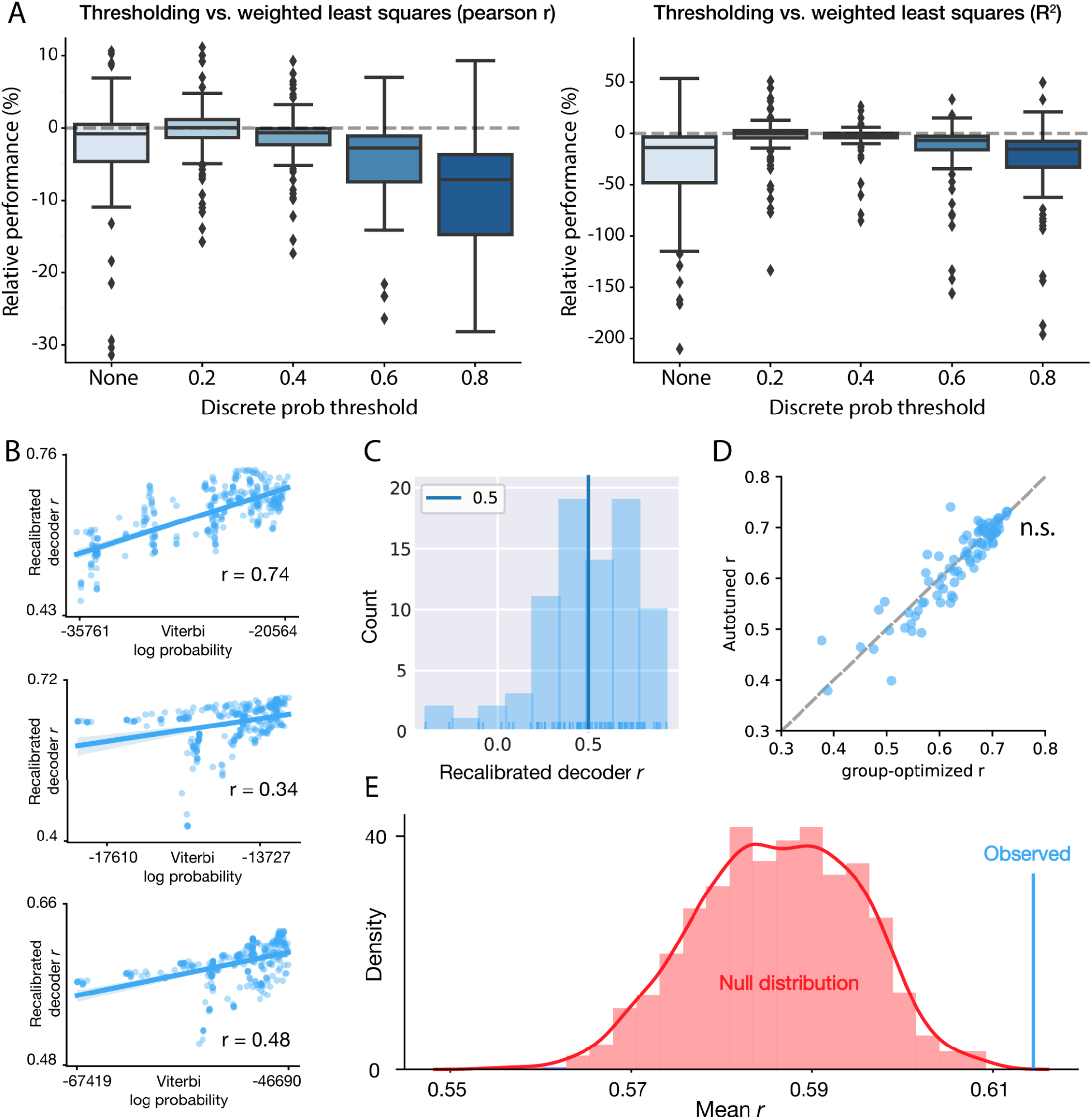
Supplementary PRI-T analyses. **(A)** PRI-T performance change using a discrete cutoff in place of weighted least squares (% relative to weighted strategy). “None” corresponds to case where all timepoints are included and equally weighted. Left: relative change in Pearson *r* (medians: −0.8%, 0.1%, −0.6%, −2.7%, −7.1%). Right: relative change in R^2^ scores (medians: −13.7%, −0.4%, −1.3%, −6.7%, −15.2%). **(B)** Example comparisons of viterbi probabilities for different hyperparameter sets against decoder performance when using those values for PRI-T. (**C**) Distribution of correlations from (B) taken across all session-pairs. Most session-pairs have a positive correlation between the two. (**D**) Comparison of recalibration strategies: performance using autotuning is slightly worse than a standard grid search (although not statistically significant, shuffle test). (**E**) Autotuning improves performance beyond expected by randomly picking values (shuffle test).

**Supplementary figure 6.**
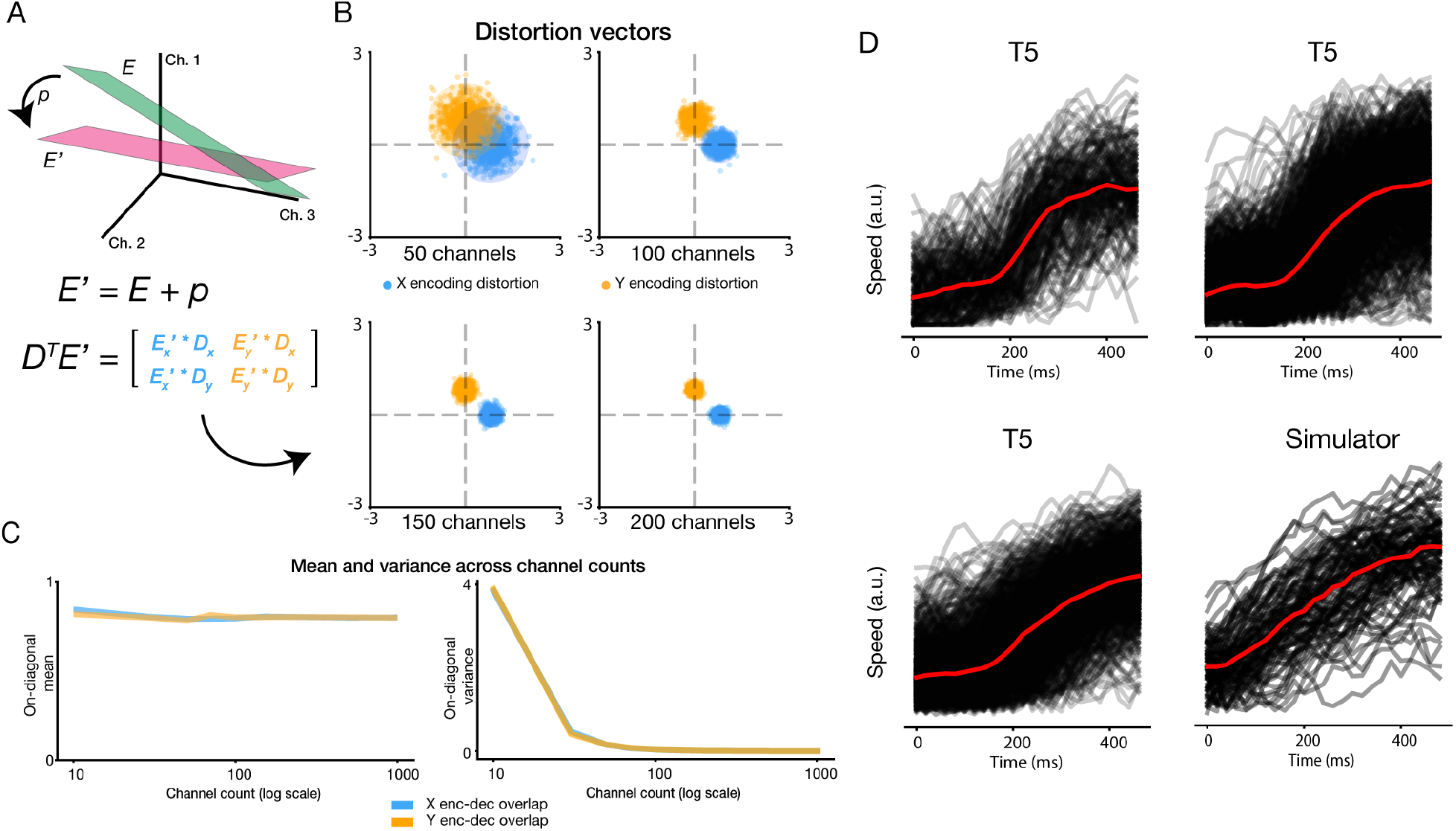
High channel count systems and simulator. **(A)** Problem overview: on each new session, the encoding space E is shifted by some random nonstationarity p, yielding the new subspace E’. The effect of this shift on a decoder D can be summarized via a distortion matrix. In an ideal setting with no neural instabilities, this is simply the identity matrix. **(B)** Simulated distortion matrices for different channel counts, plotted as columnwise vectors. The means of these distributions remain constant but their variance diminishes as channel count increases. **(C)** left: mean X and Y-subspace overlap across channel counts. Right: variance of overlap across channel counts. **(D)** Example cursor speeds, aligned to trial starts, during real and simulated closed-loop BCI use. Red lines denote median across trials.

**Supplementary figure 7.**
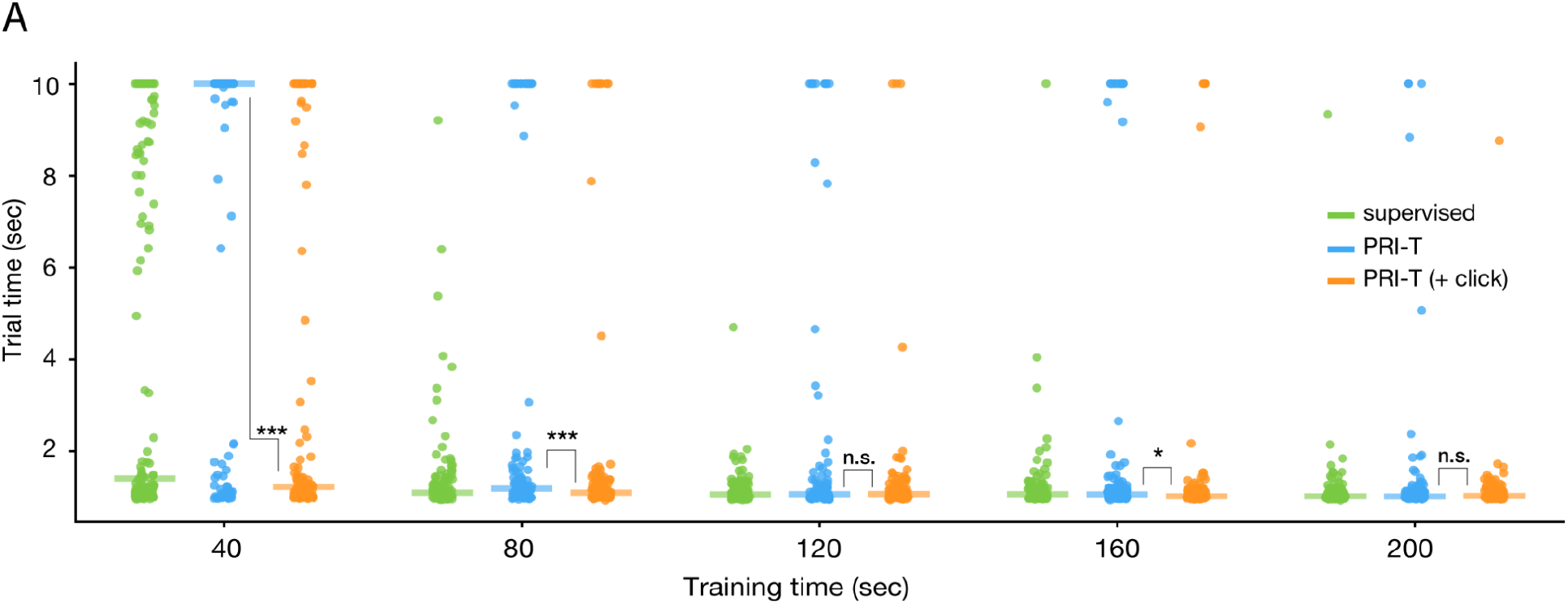
Adding click to PRI-T. **(A)** Simulated data efficiency comparison of PRI-T (orange) and PRI-T with an additional click signal integrated into its observation model. Average trial times across 200 independent runs (per method and size) are measured at 30 days out.

## Methods

Research sessions were conducted with volunteer participants enrolled in the BrainGate2 pilot clinical trial (ClinicalTrials.gov Identifier: NCT00912041). The trial is approved by the U.S. Food and Drug Administration under an Investigational Device Exemption (Caution: Investigational device. Limited by Federal law to investigational use) and the Institutional Review Boards of Stanford University Medical Center (protocol #20804), Brown University (#0809992560), and Massachusetts General Brigham(#2009000505).

Participant T5 is a right-handed man who was 69 years old at the time of the study. He was diagnosed with C4 AIS-C spinal cord injury eleven years prior to this study. T5 is able to speak and move his head, and has residual movement of his left bicep as well as trace movement in most muscle groups. T5 gave informed consent for this research and associated publications.

If a p-value is less than 0.05, it is flagged with one star (*). If a p-value is less than 0.01, it is flagged with 2 stars (**). If a p-value is less than 0.001, it is flagged with three stars (***).

### Experimental setup and neural recordings

We obtained 73 separate sessions worth of data from participant T5 spanning five years (2016-2021). While some of these data have been previously reported ^3,44^, many sessions are from monthly cursor tasks intended to monitor long-term device viability. These sessions comprise multiple tasks (“radial-8”, “grid”, and “random”, Fig. 1A) but all share a 2D cursor-to-target acquisition structure. In radial-8 and grid tasks, the screen contains a set of equally-sized targets to select from whereas the random task contains a single, often variably-sized target. The “goal” or active target in these tasks is often predetermined in advance (radial-8 and random) but these data also include free response settings (e.g. from ^3^).

From T5’s historical cursor control data, we selected sessions with ≥ 2 blocks of closed-loop cursor control recordings each. Spikes were identified based on threshold crossings at −4.5 x RMS. The resulting neural time series from each session were downsampled to 1000 Hz and then binned in non-overlapping, 20 ms bins to generate firing rate estimates. These estimates were further smoothed with a causal half-gaussian filter (*σ* = 40 ms) to denoise signals, resulting in a 192-dimensional vector 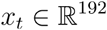 of population firing rates at each timestep *t*.

For all analyses aside from cosine tuning models (below), we also performed blockwise mean subtraction to center firing rates.

### Cosine tuning models

Cosine tuning curves ^45^ were fit to each channel by regressing the instantaneous cursor angle *θ_t_* (with respect to the target) against that channel’s firing rate *x_t_*. Cosine tuning is a standard model in the field that captures directional tuning, which has been shown to be a large signal in motor cortex ^45–47^. The standard cosine tuning model *x_t_* = *b*_0_ + *αcos*(*θ_p_* - *θ_t_*), where *θ_p_* is the preferred direction (PD) and *θ_t_* is the instantaneous velocity angle, is equivalent to a linear model of activity, *x_t_* = *b*_0_ + *b*_1_*cos*(*θ_t_*) + *b*_2_*sin*(*θ_t_*) where *b_1_* = *αcos*(*θ_p_*) and *b_2_* = *αsin*(*θ_p_*) We hence used standard least-squares to fit models by regressing neural firing rates against the ground-truth (unit) displacement vector.

Ground-truth firing rates plots (Fig. 1B) are obtained by first binning observed cursor angles into 15 evenly spaced, non-overlapping bins from 0 to 2π radians. The corresponding firing rates at each timestep are then averaged for each trial x bin x channel. We then generated bootstrapped mean estimates by sampling with replacement across trials to obtain 95% confidence intervals for each channel and bin angle. Trials with < 10 timepoints (200 ms) in a given bin were excluded from the sampling procedure.

### Measuring drift in T5’s recordings

For the offline analyses in Fig. 1, we fit and evaluated ridge regression models within each session by regressing the instantaneous displacement vector (*y_t_* = *g_t_* - *p_t_*), where *g_t_* is the target location at timestep *t* and *P_t_* is the cursor position) against population firing rates in a 50:50 train:test split across blocks. We swept L2 regularization strengths using 5-fold cross-validation and then retrained models on the full training set using the optimal value. We then measured performance using the Coefficient of Determination (R^2) on the test set.

### Bias

To measure bias across time from old decoders, we used a normalized bias index that takes into account the within-day signal-to-noise ratio (SNR). Specifically, we model a decoder’s outputs 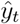 on the test set as a noisy linear readout of the true (unit-) displacement vector *y_t_* i.e. 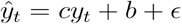 where *c* is a constant and other terms are 2D vectors. In this equation, *ε* is noise, *b* is the degree of bias in the signal, and *c* measures signal since it reflects the degree to which decoder outputs lay along the ground-truth displacement vector (after centering). We fit this equation with least-squares. If we now take the ratio ∥*b*∥/^*c*^, we get a normalized measure that takes into account both the bias on a given day as well as the tuning strength. This enables us to differentiate between days where bias is equivalent in magnitude but different in impact due to varying SNR (e.g. a high SNR system, where *c* is large, will “drown out” a rather small bias term). In practice, we fit this model to timepoints where the cursor is far from the target (> 300 pixels) to create a more interpretable final number; by measuring the decoder output far from the target, we obtain a value of *c* that reflects maximal “neural push”. A bias index above 1 therefore indicates that the decoder bias is so strong that even high-velocity signals are drowned out by the bias magnitude.

As we are modeling outputs during closed-loop control, one confounder is the possibility of within-session bias in the ground-truth data. This is problematic because an unbiased, high SNR decoder would have a large bias index in such a case despite being well-calibrated. We adjust for this by measuring the decoder’s output bias after removing the empirical bias *b*_0_ and obtain a final formula ∥*b* - *b*_0_∥/*c*.

When plotting bias values in Fig. 1E, we cap the y-axis upper limit at 10 to better highlight trends. Similarly, in the inset we cap x and y-axes upper limits at 1.2 for clarity (one outlier session had sudden, brief electrical noise that massively inflated the bias estimate for that day).

### Tuning similarity

To measure tuning similarity, we examined the cosine angle between the weights of ridge regression models after collapsing across X and Y dimension coefficients. For instance, for decoder weight matrices 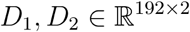, we solve for:

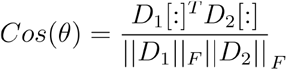

Where ∥·∥_*F*_ is the Frobenius norm. A value of 1 indicates complete overlap in the readout dimensions whereas a value of 0 indicates that the planes are orthogonal. For within-session drift we trained a decoder on the test set, while following the same cross-validated regularization procedure as earlier, and took the cosine angle of its weights against the training set decoder.

### High channel count systems

Here we explain the motivation for simulating a consistent, low-variance regime of neural changes across time, which is what would likely happen in a commercially available, high channel-count iBCI system.

We consider a simple encoding system where the firing rate vector *x_t_* across *k* channels is a noisy linear function of the intended velocity signal *v_t_*, i.e. *x_t_* ≈ *E^T^v_t_*, and representational drift occurs via random walk noise across days that preserves the tuning weights norm, i.e. 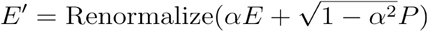 where *P* ~ *N*(0,*σI*) and with the same column norms as *E*, and *α*<1. As the encoding subspace *E’* drifts away from *E* through compounding noise, we might ask how the decoding weights D align with this new subspace.

We can compactly represent this information with a *distortion matrix D^T^E’*, which holds the X and Y-subspace overlap on the diagonal and “collisions” between the X-encoding and Y-decoding subspace on the off-diagonal (and vice versa). As the number of channels *k* scales, we find that the variance of these terms drops off like 1/k and *D^T^E’* → *αI* using a rough proof (see below) as well as simulations (S6A-C). This finding leverages the observation that high-dimensional random vectors are nearly orthogonal to each other; as the number of channels increases, the random walk drift becomes less prominent within the task subspace.

### Proof sketch

Suppose our recordings measure from *k* neurons with an associated encoding matrix 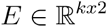, where row *E*_*i*,:_ contains neuron *i*’s x and y-velocity tuning coefficients. Denote the columns *E_x_*,*E_y_*, which are linear subspaces within which the population encoding of the velocity signals lie. We further assume that ∥*E_x_*∥ = ∥*E_y_*∥ and *E_x_* ⊥ *E_y_* and we then construct a decoder *D* = *E_T_*/∥*E_x_*∥^2^. The former two assumptions say that the encoding of horizontal and vertical movements should have the same SNR and be encoded in separate subspaces. The latter assumption supposes that we have sufficient data, and is in fact the OLE solution as the number of channels with uniformly distributed PDs approaches infinity^48^.

Now suppose that this population encoding drifts according to a decay model 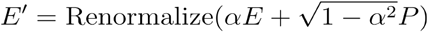 where *α* is a “shrinkage factor” reflecting diminished tuning in the original subspace and *P_i,j_* ~ *N*(0, *σ*_2_) is some random new component to the tuning. Then we have that:

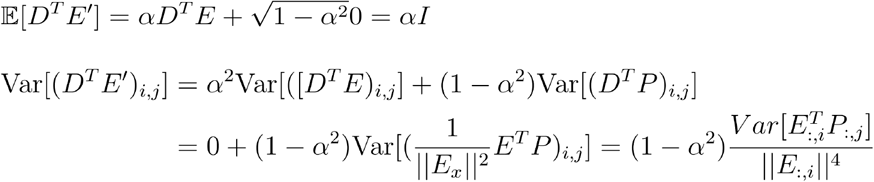

We note that 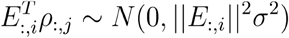, as the sum of independently distributed gaussians is gaussian with means and variances adding, and hence obtain a simplified form for the variance:

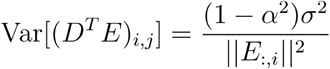

Note that, as the number of channels grows, the denominator also increases on average and implies that variance diminishes with increasingly high channel counts. If we suppose that 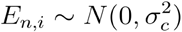 (for some neuron *n*) then 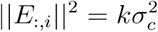 on average as 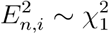. So we find that, with all else equal, the variance falls off like *1/k* for increasingly high channel count systems.

### Simulating distortion matrices

We simulated random tuning shifts 1000 times for different channel counts. On each iteration, we randomly sampled PDs for each channel from the unit sphere then scaled the resulting population tuning columns to have unit norm; we take the transpose of this matrix as the decoder D. We then generate an IID random perturbation vector *P* (same sampling as PD weights, and same norm as E) and average its weights together with the original weights *E*, followed by column renormalization, via 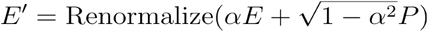. We used *α* = 0.8 in practice for the results in S6. Finally, we measure *D^T^E’* to obtain the overlap between the original decoder and the new encoding subspace.

### PRI-T

We model the cursor position *P_t_* and velocity *v_t_* (collectively, observations *O_t_*) at some timestep *t* as a reflection of the target position *H_t_* using a hidden Markov model (HMM). This has three components: 1) a description *P*(*H_t_*|*H*_*t*-1_)of how the target location evolves overtime 2) a posterior distribution *P*(*O_t_*|*H_t_*) over the observations with respect to a proposed target location and 3) a prior probability of the target location.

To model the target evolution across time, we want to maintain generality across cursor tasks while broadly capturing task structure to support PRI-T’s performance. Additionally, we require the target to behave according to discrete first-order Markov dynamics; this assumption means that the current (discrete) target location is simply a function of the previous state and helps enable fast, exact inference. We accomplish this by discretizing the screen into a *N* × *N* grid and modeling the probability of moving from one grid position to another via a matrix:

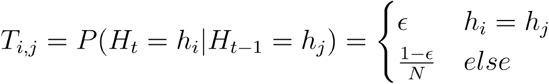

This expression says that the target has some probability *ε* of remaining in the current location and a uniform probability of transitioning to any other location. The latter decision provides task generality as we don’t assume knowledge of how the target location varies in any detail; for specific tasks such as keyboard typing, one can leverage letter bigram probabilities here to improve within-task performance although we opt for a general approach. We use *ε* = 0.999 to reflect the timescale on which the HMM operates; trials are on the order of a second whereas timesteps are 20 milliseconds. In summary, the distribution encodes a few assumptions: 1) the current target location is just a function of the previous target location 2) the target is likely to remain in its current spot for a while and 3) all other possible spots are equally probable.

To model the observed cursor state as a function of target location, we consider a Von Mises distribution over cursor velocity angles. The intuition here is that the cursor angle *θ_vt_* with respect to the current target should be concentrated around 0 degrees. This provides an initial posterior of the form *P*(*O_t_*|*H_t_*) = *VonMises*(*θ_vt_*; *κ*) where *κ* controls the concentration of the distribution around its mean and reflects the noise in the decoder’s angular outputs. We improve upon this initial description by noting that the variability of the cursor’s angle with respect to the target will increase when the cursor is near the target, as small changes in position cause large changes in angle. To accommodate this effect, we parametrize *κ* as a function of the cursor-to-target distance *d_Ht_* = ∥*P_t_* - *H_t_*∥:

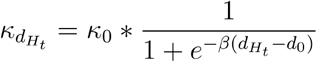

Here we weight an initial kappa value *κ*_0_ by a logistic function; at large distances the effective *κ* is close to this value. But at smaller distances kappa is closer to 0; this causes a higher variance in the Von Mises distribution which means that noisier velocity angles are likely when near a target. The exponent and midpoint variables *β* and *d*_0_ are found via an exhaustive grid search (S1A). Our final posterior is then:

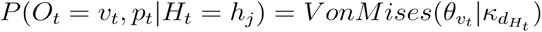

Finally, we require a prior probability over the possible target states. This distribution *P*(*H_t_*) reflects our *a priori* belief about the target location and can be used to encode task-specific regularities. For instance, if one wanted to optimize for keyboard typing this could be the empirical distribution of starting letters across all words. Like earlier, we forgo these sorts of optimizations to ensure generality across cursor tasks and instead use a uniform prior across all possible locations.

During hyperparameter sweeps, we found that using weighted least squares for incorporating timepoints based on how confident the model is about the target location was superior to using all timepoints equally or applying a discrete cutoff (Fig. S5A). We also saw that PRI-T’s Viterbi probabilities given by different hyperparameter settings were correlated with the resultant decoders’ offline performance on a new day (Fig. S5B,C; mean *r* = 0.50), meaning that hyperparameters which appeared to offer a better explanation of the data also tended to train better decoders. We therefore explored an automated tuning approach whereby the hyperparameters which maximized the Viterbi probability were selected. This approach was slightly worse than the full optimization (Fig. S5D, Δ*r* = −0.016 = 0.036; mean±SD), although not statistically significant when restricted to independent samples (p = 0.20, Wilcoxon signed-rank test), and performed better than expected by chance (S5E, p = 0.001, shuffle test).

### Click Integration

We can integrate an additional click decoder’s outputs through an indicator variable *c_t_*, which records whether or not a click occurred. To model this probabilistically, we suppose that clicks have a higher likelihood of occurring when the cursor is near the target:

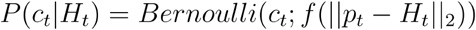

Where *f*(·) is fit to the empirical click likelihood as a function of target distance from historical sessions. We then multiply this probability with the Von Mises probability, yielding the overall posterior probability over the observations *O_t_* = {*p_t_*,*v_t_*,*c_t_*}. For the simulation in S7, we set *P_Bernoulli_* = *f*(*x*) = {1 if *x* < 0.075 else 0}, as only a single target is active during simulated control (and hence all registered clicks are valid).

### Inference

Using the HMM structure, we can then infer the most likely target sequence given the data *P*(*H*_1_,…, *H_n_*|*O*_1_,…, *O_n_*). Note that this expression is an inversion of the posterior distribution earlier, as here we now measure the likelihood of the *target locations* given the observed data. To do so, we can perform exact inference for the most likely sequence of target locations using the Viterbi search algorithm ^49^. This algorithm’s complexity is linear in sequence length, allowing for relatively fast computation. We also make use of the occupation probabilities during inference; these are the marginal probabilities of the target being in a given state at a given timestep (given the observed data). These values are obtained via the forward-backward algorithm (also linear in sequence length), and used to weight the Viterbi labels. In practice, we use the square of the maximal occupation probability at a given timestep as a weight, *max_h_P*(*H_t_* = *h*|*O*_1_,…, *O_n_*). This latter sequence of probabilities can yield sequences that differ from the Viterbi sequence but, during initial testing, noticeably improved performance.

### RTI

We implemented RTI as described in ^5^ for closed-loop simulation analyses. For each registered selection, we take a time window of length *L* (“look back” in hyperparameter sweeps) prior to that click and assign the cursor position during the selection as the intended target. The cursor-to-(inferred) target displacement signal is then regressed against neural activity from these time windows as well.

As in ^5^, we apply several heuristics to improve the performance of this retrospective labeling. First, if the labeling window extends into a previous labeling window associated with a different selection, we avoid overwriting those overlapping timesteps. Second, we remove timesteps from the window when the cursor is too close in space (“minimum distance” threshold in hyperparameter sweeps) or time (“minimum time”) to the inferred target. Finally, we remove timesteps where the distance function derivative is nonnegative i.e. when the cursor isn’t actively moving closer to the inferred target.

### FA stabilization

In ^6^, the authors use a low-rank matrix decomposition of neural activity, *X* = *LF* + *ϵ* where *L* ∈ *R^d×k^* is the loadings matrix and *F* ∈ *R^k×n^* is a low-dimensional representation of the data. Here *d* is the dimensionality of the neural recordings, *k* is the latent space dimensionality, and *n* is the number of datapoints used. This decomposition is performed via Factor Analysis in our offline results and via PCA in our simulations; during initial experiments we saw that PCA tended to provide better results for FA stabilization than FA itself. A velocity decoder is then trained in the subspace *L* by regressing velocity against the timepoints *F*_:*t*_. To recalibrate across days, a Procrustes alignment realigns the loadings matrices and *L*^(1)^ and *L*^(2)^ i.e. finding a matrix O such that *O* = *argmin*_*O*^T^*O*=1_∥*L*^(1)^ - *OL*^(2)^∥.

To make the Procrustes alignment tractable, a channel selection algorithm constrains the optimization to a subset *S* of “stable” channels ^6^. This selection algorithm uses an initial minimal stability threshold (“threshold” in hyperparameter sweeps) to prune noisy channels which have a L2 difference between their loadings weights greater than this value. The algorithm then iteratively prunes remaining channels until a requested number of stable channels remains (“B” in hyperparameter sweeps). The full alignment is then 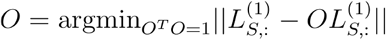 where S indexes stable channels.

### Daisy-chaining

As drift compounds, fewer stable channels exist and this mapping becomes increasingly difficult. We avoid this problem by dividing the alignment into a series of subproblems: aligning between consecutive days when drift is small. To do so, we first initialize the stabilizer as in ^6^ and obtain 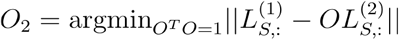. On a new day, we then repeat the channel selection algorithm and procrustes alignment, obtaining 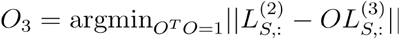. We can then compose these transformations to obtain *O*:= *O*_2_*O*_3_. This process is run again for each subsequent day. In simulation, this delivered a nearly 4x reduction in trial times from 8.73±2.52 to 2.36±2.52 seconds at the 60 day mark in simulation (Fig. 4D).

### ADAN

During initial within-day model training, we found that the original ADAN architecture tended to massively overfit to behavioral task data. We therefore introduced two changes to combat this problem. First, we swapped out the RNN task predictor for an FCN with a 10-dimensional hidden layer. Second, we introduced multiple data augmentation strategies to improve noise robustness and combat overfitting. This was accomplished by adding constant channel offsets, per-timestep white noise, and random walk noise to each minibatch:

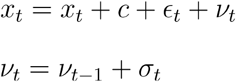

where *c*, *ϵ_t_*, *σ_t_* ~ *N*(0,*q*). We used q = 3, 14, 0.9 respectively. Models were trained for 50 epochs with an Adam optimizer (learning rate = 1e-3) and minibatch size of 64.

Across-day alignment hyperparameters were tuned through a grid search (Fig. S2) over batch size, generator learning rate, and discriminator learning rate, based on results from ^27^ demonstrating performance being especially sensitive to these variables. Because of compute requirements (27 parameter sets x ~1 hr of GPU wall-clock time = 27 hours per session-pair), we used a subset of 30 session-pairs for sweeps. During initial analyses, we observed that results tended not to converge for small epoch sizes; we consequently used 200 epochs as in ^28^. We otherwise use default hyperparameters.

### Offline hyperparameter sweeps

We performed exhaustive grid searches for PRI-T, FA stabilization, and ADAN over pairs of sessions. Since all methods tended to fail on sessions without sufficient tuning changes (Fig. 3C), we opted to subselect session-pairs where a mean recalibration baseline satisfied a threshold of *r*_mean recal_ > 0.15^0.5^. Hyperparameters that maximized the mean performance across session-pairs were then selected.

Below we reproduce the hyperparameter values tested and selected for all three methods:

#### PRI-T

| Hyperparameter | Values swept | Optimal value |
| --- | --- | --- |
| Base kappa | 0.5, 1, 2, 4, 6, 8, 10 | 2 |
| Logistic inflection | 0.1, 10, 30, 50, 70, 100, 200, 400 | 70 |
| Logistic exponent | 0.01, 0.1, 0.5, 1, 2, 4 | 0.5 |

#### FA Stabilization

| Hyperparameter | Values swept | Optimal value |
| --- | --- | --- |
| Latent dimensionality | 2, 4, 6, 8, 10, 12 | 6 |
| B | 80, 100, 120, 140, 160, 180, 190 | 160 |
| Threshold | 1e-5, 5e-5, 1e-4, 5e-4, 1e-3, 5e-3, 0.01, 0.05, 0.1 | 0.01 |

#### ADAN

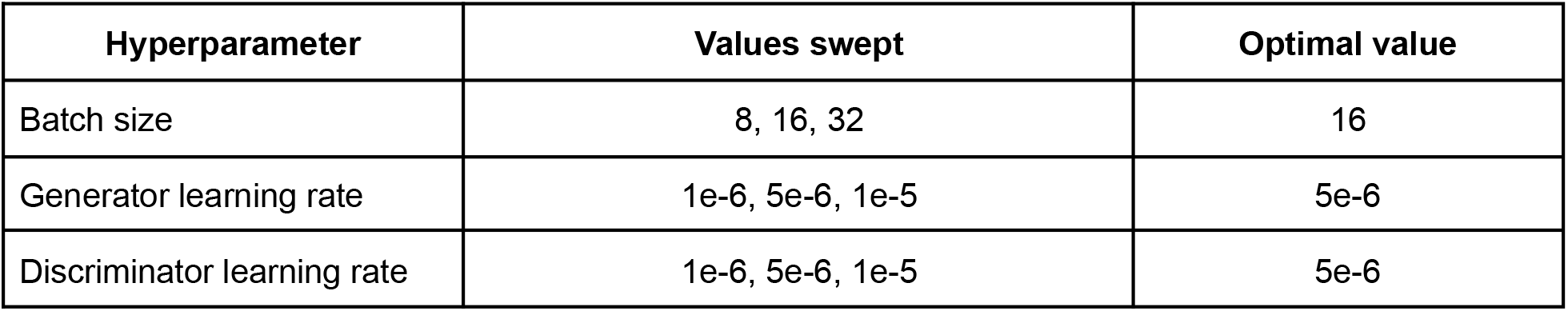

### Offline comparisons

We generated offline datasets by creating pairs of cursor sessions, with the first item in the pair being a reference session for decoder training and the second being an evaluation session. These data were processed as before (causal gaussian smoothing with *σ* = 40 ms, 20 millisecond binned firing rates, blockwise mean subtraction). Within each day, we performed a 50:50 train/test split across blocks. Sessions with at least two 2 blocks of cursor control data were used.

For examining recalibration, decoders were recalibrated on a new session’s train set and evaluated on the test set. We measured both Pearson correlation across both outputs (*r*) as well as the coefficient of determination (R^2, uniformly averaging over X and Y dimensions). While *r* and R^2 scores showed a strong, monotonic relationship, we opted for *r* as it reflects the strength of the relationship between decoder outputs and ground-truth after accounting for a difference in means and scaling. The former is modified by online mean subtraction algorithms during closed-loop control and the latter is often dealt with via simple gain scaling procedures that are also modified for optimal control. In Fig. 3C, we plot *r*^2^, which is an upper bound on *R*^2^ assuming gain and bias are controlled for.

In Fig. 3B, we took care to subselect days where mean recalibration had high performance. This reduces the within-approach variance by removing far apart sessions where all methods tend to fail. Despite thresholding pairs based on mean recalibration (*r*_mean recal_ > 0.15^0.5^), all methods nonetheless outperformed mean recalibration. We apply this same thresholding to pairwise comparisons in Fig. S3 and S4 below, as well as removal of an outlier day with sudden electrical noise.

We also looked more directly at pairwise comparisons (Fig. S3). All trends were roughly consistent with the above initial analyses. PRI-T, ADAN, and FA stabilization outperformed mean recalibration (>75% of cases for all three, Fig. 4A) and with no major differences in median performance; among pairwise comparisons of the three, the largest improvement was a median 1.7% increase in scores using PRI-T over ADAN (Fig. S3B). Using a combined approach was slightly worse than supervised recalibration (4% improvement with supervised, Fig. S3C) but better than PRI-T or FA stabilization alone (>85% of cases). Similar results were obtained when displaying R^2^ as well (S4).

### Closed-loop simulator

The closed-loop simulator here is a variant of the PLM simulator described in ^38^, but with an additional neural tuning model and simplified noise. A control policy *f_targ_* generates a command signal *c_t_* at each 20 millisecond timestep in response to delayed visual feedback:

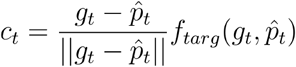

where *g_t_* is the current target location. The simulated user’s internal estimate of the cursor position 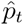 is obtained by forward-integrating from the ground-truth cursor position 200 milliseconds ago, using the previous command signals and ground-truth smoothing and gain ^38^.

This command is encoded across *k* channels with linear tuning to its x and y components (uniformly sampled preferred directions) to simulate neural activity *x_t_* at timestep *t*. Neural activity is then corrupted by independent, mean-zero gaussian noise (*σ* = 0.3) and linearly decoded at each timepoint, yielding a noisy velocity command signal *d_t_*:

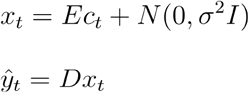

This signal is exponentially smoothed with the prior velocity (*α* = 0.94) and scaled by a fixed gain *β* (determined via sweeps, described below) before updating the cursor state.

### Matching simulator and participant encoding strength

For comparing SNR between T5 and our simulation, we use a SNR metric that models offline decoder outputs 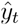 as a scaled, noisy readout of the true point-at-target vector 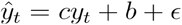 where *b* is a constant bias term, *c* is a scalar reflecting the overlap between true and intended command signals, and *ϵ* ~ *N*(0, *σ*^2^). We can then define the SNR of a single recording session as 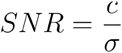. This metric is highly correlated with offline *R^2^* (r = 0.73 and r = 0.92 for experimental and simulated, Fig. 4B insets) but is insensitive to gain miscalibration on a test block as the *a* term absorbs constant gain differences. We fit this model using offline linear decoder outputs, while avoiding timepoints that were either 1) shortly after a new trial began (within 200 ms for T5, within 140 ms for simulator) or 2) close to the target (< 300 pixels for T5, < 0.3 units for simulator).

We then changed the tuning strength of our simulated channels to match T5 by changing the overall magnitude of the tuning coefficients (the “population PD norm”) across channels, ∥*E_x_*∥ where *E_x_* = [*e*_1_,…, *e*_192_]^*T*^, to align with the SNR values obtained empirically from T5. When plotting the simulated population PD norm against the associated SNR across simulation runs, we saw a tight correlation (*r* = 0.99). This meant we were able to adjust the simulator’s SNR distribution by simply finetuning the population PD norm distribution. To find an appropriate PD norm distribution, we then fit a gaussian mixture model (n_gaussians = 2) to T5’s SNR using expectation-maximization. Simply rescaling the value of this distribution then provides the correct PD norm distribution in the simulator. During simulator use, we randomly sampled from the model, followed by scaling and bias to adjust the output to the corresponding magnitude value:

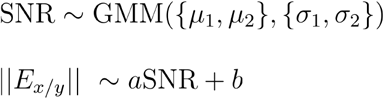

### Matching simulator and participant encoding drift

To examine encoding drift across time in our participant recordings, we measured the cosine angle difference *cos*(*θ_i,j_*) of the encoding weights for different pairs of sessions *i* and *j* (and separately for x and y-subspaces within these pairs). We then fit exponential decay models that suppose that *cos*(*θ_i,j_*) decays by some constant multiple across time i.e. cos(*θ_i,j,k_*) = *bα^k^* where *b* is a constant and set to 1 in practice; this latter assumption supposes that the within-day encoding is stable. We can then estimate *α* by solving the least-squares problem cos(*θ_i,j+k_*) = *α*^k^ in log-space, log(cos(*θ_i,i+k_*)) = *k*log(*α*), which amounts to linear regression. As the logarithm is undefined for negative values, we mask data points where the cosine angle value is less than zero in the optimization. The resulting coefficient is the logarithm of our decay parameter; exponentiating this value yields the decay estimates *α* = 0.913 and *α* = 0.925 in Fig. 4C.

Next we would like to instantiate the same drift rate in our simulated units. To do so, let’s again consider the drift model 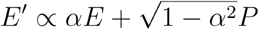. With a few simple assumptions, the shrinkage parameter *α* here aligns with the cosine-angle difference decay parameter described above. To see this, consider the cosine angle difference:

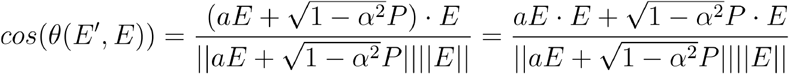

Now we make two simplifying assumptions: 1) *P* is orthogonal to E and 2) that the magnitude of E’ is roughly equal to E. The first assumption is relatively safe for high-dimensional recordings (random vectors are nearly orthogonal in high dimensions) and the latter is equivalent to saying that SNR is roughly equivalent across days. This yields:

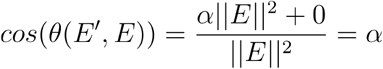

In practice we use an explicitly orthogonal *P* (as this mimics the anticipated task setting of recalibration in the high-dimensional channel count regime) but vary the tuning strength of the simulator, which violates the second assumption. However, the approximation still holds well as we obtain an empirical cosine angle decay of *α* = 0.909 when using *α* = 0.910 for our shrinkage parameter.

### Performance sweeps

We simulated two months of data for each recalibration approach by first building an initial linear decoder on day 0. All decoders used 20 seconds of open-loop data and a fixed smoothing coefficient of 0.94. Optimal gain was determined by sweeping across ten, evenly spaced values from 0.1 to 2.5 and measuring closed-loop trial times (200 second blocks).

We then applied different recalibration schemes while slowly modifying neural population activity. On each new day, we update the encoding matrix E with an orthogonal P via 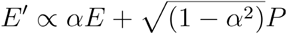, where *α* = 0.91. We then rescale the column magnitudes by sampling from the GMM fit to T5’s SNR distribution (see above). Next we apply the previous day’s decoder and obtain 200 seconds of closed-loop control data. This block is used for recalibration, followed again by an optimal gain sweep, and the resulting decoder is tested once more on a new block to obtain trial time measures. The decoder is then saved and redeployed again the following day after following the above steps once more. In practice, we independently initialize and update the encoding matrix for each run to maintain statistical independence across simulations. Each method is simulated in 200 such independent runs.

For dataset size sweeps, we reduced the length of the recalibration block while measuring average trial times on a fixed size evaluation block lasting 200 seconds.

### Simulator hyperparameter sweeps

We tested three unsupervised recalibration methods in simulation (PRI-T, FA stabilization, and RTI) as well as a simple gain recalibration baseline where decoder weights were fixed and only the gain was optimized.

For all methods, we simulated 30 days of unsupervised recalibration, with slowly varying neural tuning as described earlier, and applied various hyperparameter settings in a grid search fashion.

We tested each hyperparameter set with 30 independent runs. The hyperparameter sets that minimized the median trial time for each method at the 30 day point were selected for subsequent performance comparisons.

#### PRI-T

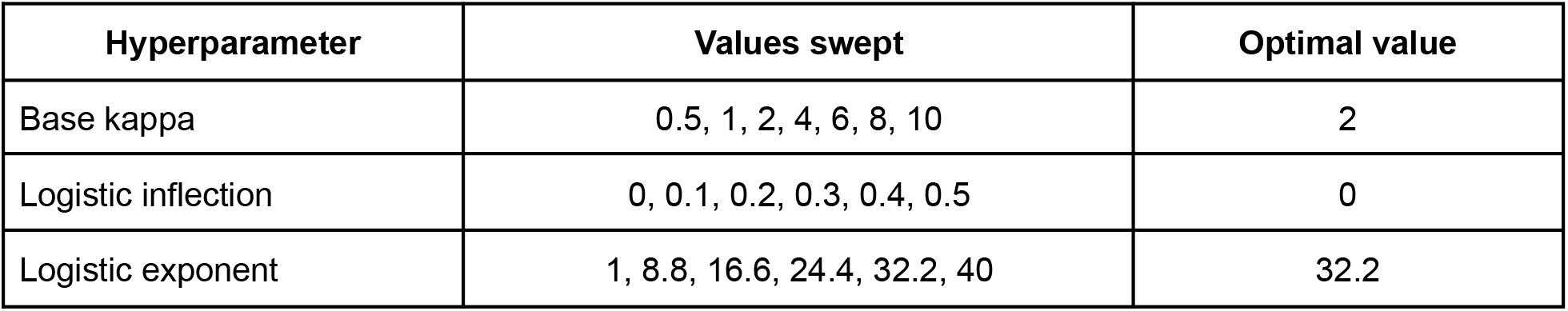

| Hyperparameter | Values swept | Optimal value |
| --- | --- | --- |
| Base kappa | 0.5, 1, 2, 4, 6, 8, 10 | 2 |
| Logistic inflection | 0, 0.1, 0.2, 0.3, 0.4, 0.5 | 0 |
| Logistic exponent | 1, 8.8, 16.6, 24.4, 32.2, 40 | 32.2 |

#### FA Stabilization (daisy-chained)

| Hyperparameter | Values swept | Optimal value |
| --- | --- | --- |
| Latent dimensionality | 2, 4, 6, 8, 10, 12 | 2 |
| B | 10, 40, 70, 100, 130, 160, 190 | 100 |
| Threshold | 0.01, 0.05, 0.1, 0.2, 0.3 | 0.05 |

#### FA stabilization (static)

We used 10 repetitions here instead of 30, as early results indicated that static FA stabilization was highly unstable relative to daisy-chaining.

| Hyperparameter | Values swept | Optimal value |
| --- | --- | --- |
| Latent dimensionality | 2, 4, 6, 8, 10, 12 | 6 |
| B | 10, 40, 70, 100, 130, 160, 190 | 190 |
| Threshold | 0.01, 0.05, 0.1, 0.2, 0.3 | 0.1 |

#### RTI

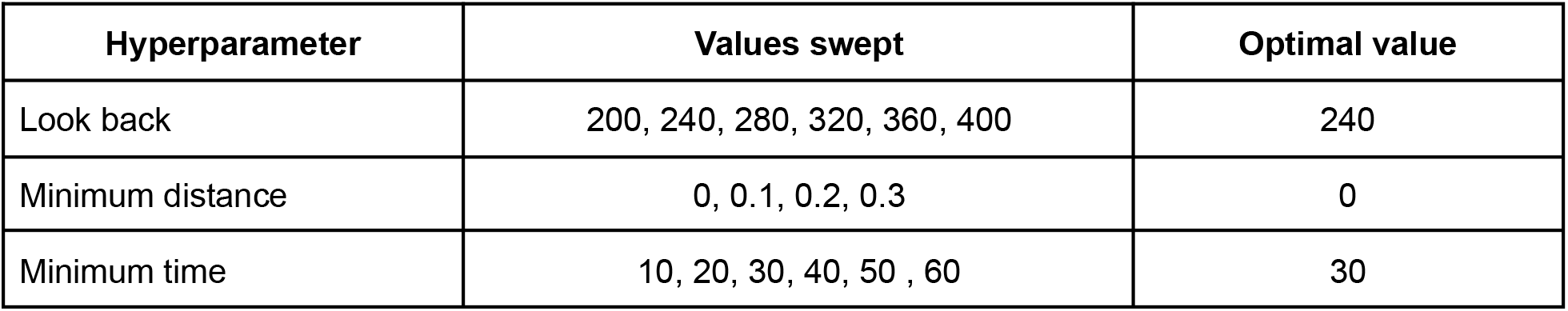

### Closed-loop evaluation

Task logic and visualization were implemented in Python using the PyQtGraph library. At the start of a trial, a variable-size target is generated randomly on the screen. The user must then move the cursor over the target and dwell for a consecutive, 500 millisecond period to select it. Trials lasting more than 10 seconds were counted as failures. Upon trial completion, audio feedback is provided to the user to indicate whether or not the trial was successful and a new target is generated immediately. OL and CL blocks were 2 and 4 minutes long, respectively.

Neural data were binned in 20 millisecond, non-overlapping bins and fed through a linear regression model *D* trained to predict the cursor-to-target displacement (defined as *g_t_* — *P_t_*, where *g_t_* is the current target position and *P_t_* is the cursor position). The raw velocity signal *v_t_* was exponentially smoothed with the running velocity average *c_t_* and scaled with a gain parameter *β* via *c_t_* = *αc_t-1_* + (1 - *α*)*βv_t_*, where *α* is the smoothing term and *β* is the gain parameter. Smoothing and gain were manually adjusted during the first session and fixed on subsequent days. We also applied bias correction (as in ^3^) to reduce the impact of mean shifts by subtracting a running estimate of the decoder bias *B_t_* from the raw velocity output. We used an adaptation rate of 0.3 for bias updates.

After a CL block was over, the running bias estimate was saved to the deployed decoder’s file in order to provide a better initial bias estimate when used later that same session (or the following session). We obtained confidence intervals for average trial times using bootstrapping with 10,000 iterations.

### Initial day

On day one, we trained an initial linear decoder usingT5’s neural activity while he engaged in an OL block. This decoder was then used to drive CL control in a subsequent block. We then built two decoders off of this CL block: a standard decoder using the full neural time series and an FA subspace decoder (n_components = 6). Both decoders were trained to predict the cursor-to-target displacement signal. The standard decoder was then used as the “stale” baseline decoder and also as the initial decoder for PRI-T. The subspace decoder, as well as the associated FA model, are used as the reference session for FA stabilization.

### Subsequent sessions

We collected an OL block at the beginning of each new session and later measured decoder performance offline (correlation of outputs with ground-truth displacement vector). We obtained 95% empirical confidence intervals by bootstrap resampling paired prediction and ground-truth timepoints 10,000 times.

To counteract strong session-to-session mean firing rate changes, we collected a 30 second rest block on each day where T5 was instructed to relax. We then updated the intercept term of all decoders such that the average decoded velocity vector during the rest block was zero, enforcing that the cursor not move when the user is not actively doing anything.

With all three approaches (fixed, FA stabilization, PRI-T), we then used the corresponding decoder from the prior session in an initial CL unsupervised recalibration block. We then recalibrated the decoder weights using the block’s data, followed by column-wise rescaling to match the initial day 0 decoder column norms (to keep the overall gain relatively constant), and then tested it in an evaluation CL block. We then repeated this process 1-2 more times depending on the amount of time available in the session. We interleaved blocks from the three methods in an A/B/C fashion to avoid biasing results from performance differences across time.

Decoders were recalibrated for each of the three methods as follows. For the mean recalibration strategy, we simply run the CL block with the fixed decoder; bias is automatically corrected for via bias correction. We then initialized bias correction on subsequent blocks using the latest bias correction state from the previous block. This latter strategy was not used with subspace stabilization or PRI-T, as decoder weights are updated after each CL block (and hence the estimated bias differs from the prior block’s estimate).

For FA stabilization, neural data were smoothed offline (causal half-gaussian filter, *σ* = 40 ms) and reduced to a 6-dimensional FA subspace, then realigned to the reference subspace (as in ^6^; B = 100, threshold = 0.01). At run-time, neural data are pushed through these mappings then decoded by a linear regression fit on day one. This amounts to a reduced-rank regression (RRR), *y_t_* = *DQEx_t_* where *E* ∈ *R*^6×192^, *Q* ∈ *O*(6), and *D*^2*x*6^. The outputs are then smoothed and debiased as described earlier.

For PRI-T, we infer pseudo-target labels then recalibrate by training a new decoder from scratch with weighted least squares.

When displaying trial times for a given session, we removed the first 60 seconds from the first block of each decoder. This burn-in period generally contains heavy bias, which is gradually removed by an adaptive bias correction algorithm.

